# Greatwall depletion from Xenopus oocytes reveals a key role of the cyclin B/CDK1-PP2A-B55 balance in the coordination of meiotic events

**DOI:** 10.1101/2025.10.03.680315

**Authors:** Sylvain Roque, Cedric Hassen Khodja, Célia Ben Choug, Suzanne Vigneron, Véronique Legros, Guillaume Chevreux, Benjamin Lacroix, Anna Castro, Thierry Lorca

## Abstract

Meiotic progression relies on maintaining a precise balance between cyclin B/CDK1 activity and the phosphatase PP2A-B55. The latter is negatively regulated by the Greatwall kinase (Gwl). In Xenopus oocytes, we show that the loss of Gwl and the subsequent hyperactivation of PP2A-B55 severely disrupt the transition from meiosis I to meiosis II. This disruption prevents phosphorylation of both Wee1/Myt1 and the APC/C complex, thereby blocking APC/C activation. As a consequence, APC/C remains inactive during the MI–MII transition, which impairs cyclin degradation and the partial CDK1 inactivation that is normally required at this stage. Additionally, the mos-MAPK-Rsk1/2 pathway fails to activate due to insufficient Mos accumulation, thereby preventing metaphase II arrest. Finally, the lack of APC/C activation during meiosis I inhibits the degradation of its inhibitor Erp1, revealing a critical feedback loop between APC/C and Erp1. Overall, our findings reveal that Gwl is a key coordinator of both meiotic divisions, acting through dual regulation of APC/C and the Mos/MAPK/Rsk1/2 pathways by modulating PP2A-B55 activity.

## INTRODUCTION

Meiosis is a specialized cell division process that reduces the chromosome number by half, generating genetically distinct haploid gametes essential for sexual reproduction. Additionally, meiosis introduces genetic diversity through the recombination and segregation of homologous chromosomes^1^. This process plays a critical role in sexual reproduction and is tightly regulated by the coordinated activity of kinases, including Protein Kinase A (PKA) and cyclin B1/B2/CDK1 (hereafter referred to as ’cyclin B/CDK1’), which orchestrate key protein phosphorylation events and ensure timely cell cycle transitions.

PKA activity is essential at early stages of meiotic division to establish a prophase I arrest ensuring that oocytes remain fully grown and developmentally competent for fertilization. This arrest is mediated by PKA via two parallel mechanisms: (1) suppressing mRNA translation of key cell cycle regulators, and (2) downregulating cyclin B/CDK1 activity through Myt1-dependent phosphorylation of CDK1 at its inhibitory residues (T14 and Y15). Next, progesterone (Pg) releases prophase I arrest by downregulating PKA, inducing the activation of Cdc25 and cyclin B/CDK1 and promoting meiotic resumption.

Oocytes then progress into anaphase I and exit this phase of meiosis by inducing a partial degradation of cyclin B1/B2. The incomplete loss of this cyclin guarantees the presence of a residual cyclin B/CDK1 activity essential for preventing an intermediate S phase from occurring between meiosis I and II and ensuring a correct first reductional division².

Finally, at the end of meiosis, oocytes accumulate Emi2/Erp1 protein, an Anaphase Promoting Complex/Cyclosome (APC/C) inhibitor that stabilizes cyclin B1/B2 keeping cyclin B/CDK1 activity high and maintaining metaphase II until fertilization. Notably, Erp1 expression starts from GVBD^2^ but its accumulation is prevented by cyclin B3/CDK1-dependent phosphorylation that promotes its ubiquitination and degradation. At the end of meiosis I, cyclin B3 is degraded by the APC/C^3,4^ and this negative modulation is lifted permitting the stabilization of Erp1 by c-Mos-MAPK-Rsk1/2-dependent phosphorylation and thereby enabling APC/C inhibition and metaphase II arrest^5^.

Besides the role of kinases, the regulation of phosphatases, and notably of PP2A-B55, is also crucial to enable a correct timing of protein phosphorylation and meiotic progression^6^. The mechanisms controlling PP2A-B55 activity have been largely studied in the context of mitotic division. In this process, PP2A-B55 is controlled by the Gwl kinase also known as MASTL (Microtubule-Associated Serine/Threonine Kinase-Like) in humans. Gwl is turned off during interphase when PP2A-B55 remains active and dephosphorylates key mitotic regulators such as Myt1, Wee1, and Cdc25, thereby preventing cyclin B/CDK1 activation, substrate phosphorylation and entry into mitosis^7–9^. At mitotic entry, Gwl kinase is turned on and phosphorylates its substrates ENSA and Arpp19 converting them into potent inhibitors of PP2A-B55 and enabling the subsequent phosphorylation of Myt1, Wee1, and Cdc25 finally activating cyclin B/CDK1^10,11^. Unlike mitosis, the role of Gwl kinase in meiotic maturation is not fully understood. In this study, we used the TRIM-Away technique to selectively degrade Gwl in Xenopus oocytes and assess its function during meiosis. Our results show that Gwl depletion leads to sustained PP2A-B55 activity, which in turn prevents phosphorylation and activation of both the APC/C complex and the Mos-MAPK-Rsk1/2 pathway. As a result, APC/C remains inactive, leading to the accumulation of cyclins B, and failure to degrade its inhibitor Erp1. Moreover, Mos fails to accumulate, preventing downstream activation of Rsk1/2 and thereby compromising metaphase II arrest. These combined defects severely impair both the transition from meiosis I to meiosis II and the establishment of metaphase II arrest. Together, our findings reveal that Gwl is essential for orchestrating the meiotic program by coordinating both APC/C activation and Mos pathway engagement through regulation of PP2A-B55.

## RESULTS

### Depletion of the Gwl kinase does not prevent meiotic resumption in Xenopus oocytes

In order to assess the role of Gwl in meiotic progression, we first checked the expression pattern of this kinase during meiotic maturation in Xenopus oocytes. As shown in Figure S1A, Gwl protein is stably expressed in oocytes at all stages including fully-grown stage VI immature oocytes (Prophase I-arrested oocytes: PI), maturing oocytes 2 hours upon Pg stimulation (GV+2h), oocytes upon Germinal Vesicle Breakdown (GVBD) and metaphase II-arrested oocytes (MII). As expected, this kinase displays a delayed electrophoretic mobility, indicative of its phosphorylation and activation, starting at GV+2h and reaching its minimal mobility at GVBD and MII oocytes. Complete Gwl phosphorylation is concomitant with Plx1 and cyclin B/CDK1 activation as indicated by Plx1 phosphorylation on T210 activatory site and CDK1 dephosphorylation of Y15 inhibitory residue. We next depleted Gwl from prophase I-arrested oocytes using the TRIM-Away approach^12^. This technique employs the TRIM21 protein, an E3 ubiquitin ligase that associates to antibody-bound targets and facilitates their degradation via the proteasome^13^. As expected, the microinjection of these oocytes with the mRNA encoding HA-tagged TRIM21 and anti-Xenopus Gwl antibodies promoted the complete depletion of Gwl, an effect that was not observed when control anti-GST instead of Gwl antibodies were used (Figure S1B). We next stimulated these oocytes with Pg and scored the presence of a white spot in the animal pole as a marker of GVBD. Both control and Gwl-depleted (ΔGwl) oocytes presented a white spot at four hours upon Pg addition although this spot was larger in the latter group of oocytes that rapidly degenerated in a few hours (Figure 1A and S1C). To confirm that this white spot was associated with meiotic resumption, we checked the phosphorylation of CDK1 on Y15, of Plx1 on T210 and of the endogenous CDK1 substrate T320 of PP1. No differences in the phosphorylation patterns of these proteins were observed between control (CT) and ΔGwl oocytes indicating that Gwl depletion enables normal cyclin B/CDK1 activation and meiotic resumption (Figure 1B).

**Figure 1.**
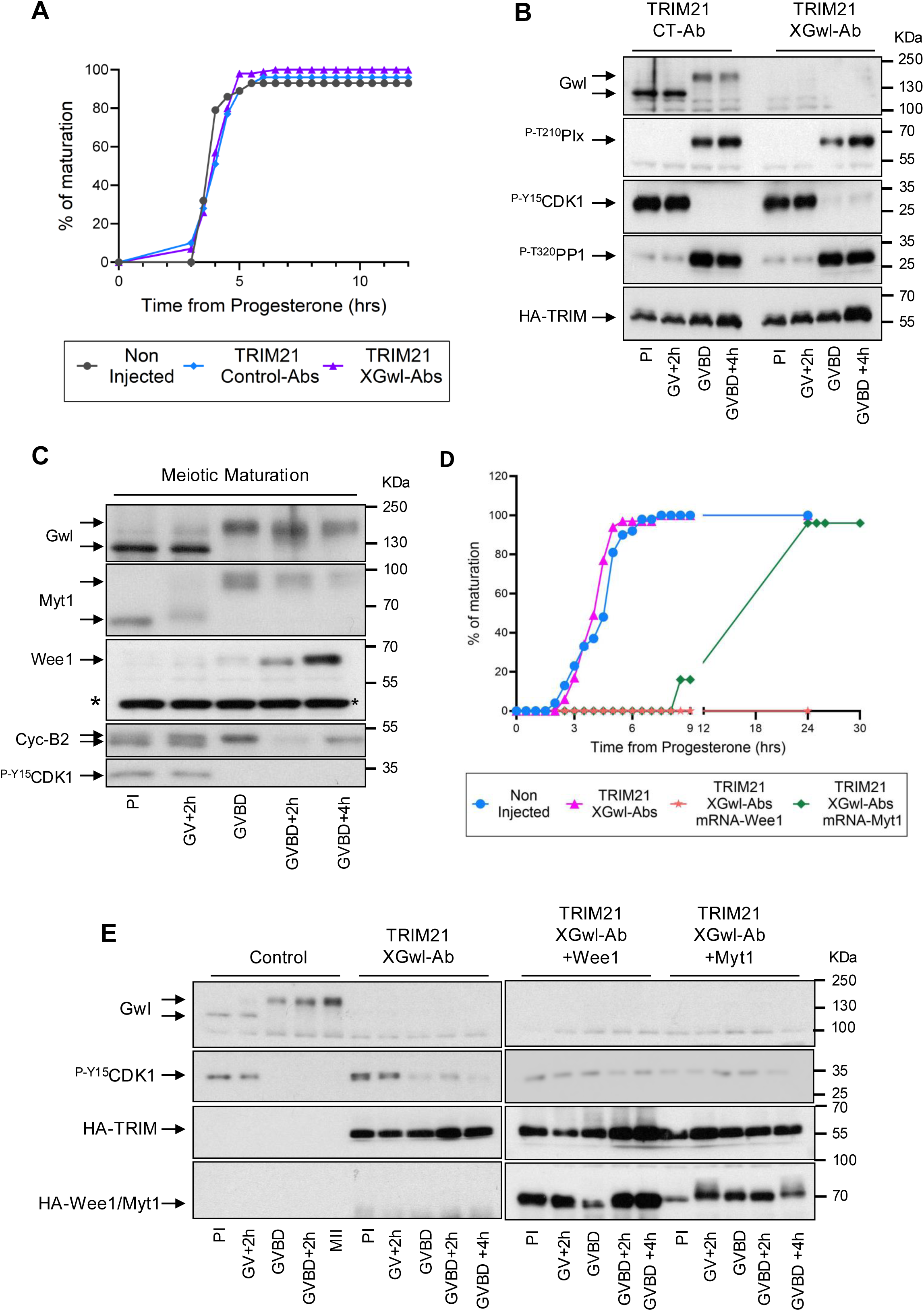
Depletion of the Gwl kinase does not prevent meiotic resumption in Xenopus oocytes. **(A)** Prophase I arrested oocytes were injected with HA-TRIM21 mRNA and either control or Xenopus anti-Gwl antibodies (XGwl-Abs). Sixteen hours later, progesterone was added and Germinal Vesicle Breakdown (GVBD, marked by the appearance of a white spot in the animal pole of the oocytes) was monitored over time. **(B)** Oocytes in (A) were recovered at prophase I (PI), two hours upon Pg addition (GV+2h), at GVBD and four hours upon GVBD (GVBD+4h) and analysed by western blot to detect Gwl and TRIM21 proteins, as well as the phosphorylation status of Plx1 (T210), the phosphatase PP1 (T320), and Cdk1 (P-Y15). **(C)** Oocytes arrested in prophase I were treated with Pg and the levels of Gwl, Wee1, Myt1 and cyclin B2 and the phosphorylation of CDK1 on Y15 were measured at the indicated timepoints. **(D)** Prophase I-arrested oocytes were either injected or not with TRIM21 mRNA and Gwl antibodies, along with Wee1 or Myt1 mRNA. Sixteen hours later, oocytes were treated with progesterone, and GVBD was monitored over time. **(E)** Oocytes in (D) were removed at the indicated time-points and analysed by Western blot to examine the levels of TRIM21, HA-Wee1, HA-Myt1, Gwl and phosphorylation of Cdk1 on Y15. Data (graphs and western blots) are representative of at least three different experiments.

Intriguingly, unlike Gwl-devoid immature oocytes, it has been shown that Gwl depletion in G2 first embryonic Xenopus egg extracts prevents cyclin B/CDK1 activation by maintaining Y15 phosphorylation of CDK1^9^. A major difference between G2 egg extracts and stage VI oocytes is that Wee1 phosphatase is not expressed in stage VI whereas it is already present in first embryonic division^14^. A putative explanation for this discrepancy could be that, because of the absence of Wee1, immature oocytes would be unable to fully phosphorylate CDK1 on Y15 in the absence of Gwl. To investigate this hypothesis, we overexpressed either Myt1 or Wee1 in stage VI oocytes and checked their effect in meiotic maturation. As previously described, while Myt1 kinase was already present in immature oocytes, we observed the expression of Wee1 in control non-treated oocytes from two hours after GVBD concomitantly with the partial drop of cyclin B2 typically observed in the oocytes between MI-MII (Figure 1C). Interestingly, the overexpression of Myt1 in ΔGwl oocytes promoted 100% of maturation albeit with a drastic delay and residual Y15 phosphorylation in CDK1 (Figure 1D and E). Conversely, Wee1 overexpression, completely blocked meiotic resumption, which suggests that the absence of Wee1 in Gwl-devoid oocytes enables meiotic progression despite the presence of hyperactive PP2A-B55 phosphatase.

### Gwl depletion induces drastic changes in the protein phosphorylation pattern of meiotic oocytes

To further characterize the role of Gwl in meiotic progression, we performed a label-free phosphoproteomic analyses in oocytes at GVBD and four hours later (GVBD+4h) and we examined the phosphorylation changes induced by Gwl loss and the subsequent reactivation of PP2A-B55. We analysed four groups, GVBD and GVBD+4h depleted (ΔGwl) or not (CT) of Gwl. For each group, we used five replicates of 10 control and 10 ΔGwl oocytes coming from different frog females (Figure 2A) and we conducted a LC-MS/MS-based phosphoproteomic analysis. Data were first analysed using a Principal Component Analyses (PCA) for the four different groups to assess factors driving phosphorylation variation. PCA outputs revealed that principal component 1 (PC1) produced two clearly separated groups corresponding to control and Gwl depleted oocytes (Figure 2B) indicating that Gwl depletion induced major phosphorylation differences, while principal component 2 (PC2) separated GVBD from GVBD+4h oocytes suggesting that developmental stage has a minimal effect on the phosphoproteome. In a second PCA, the temporal variable was removed by combining the two stages of control and ΔGwl replicates. Again, PC1 clearly separated control from ΔGwl oocytes, highlighting the strong effect of Gwl depletion in protein phosphorylation, while PC2 only captured residual variability (Figure 2C).

**Figure 2.**
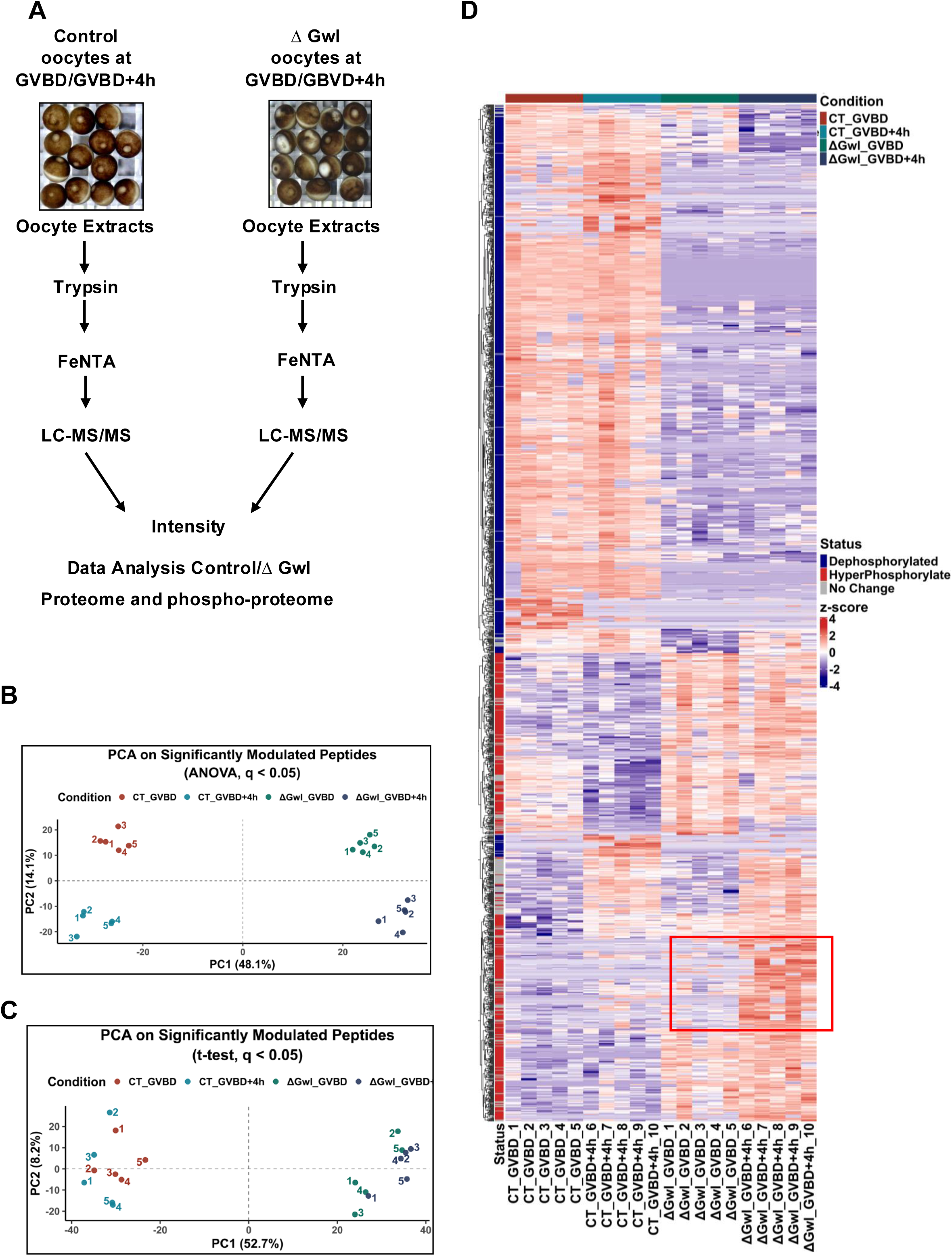
Gwl depletion induces drastic changes in the protein phosphorylation pattern of meiotic oocytes. **(A)** Scheme of the protocol used for phosphoproteomic analysis. **(B)** Principal component analysis (PCA) of significantly modulated phosphopeptides. Plot visualizing the distribution of biological replicates along the first two principal components based on phosphopeptide intensities significantly altered across conditions (ANOVA with FDR correction, q < 0.05). Four experimental conditions are represented: control GVBD (red), control GVBD+4h (blue), ΔGwl GVBD (green), and ΔGwl GVBD+4h (grey). PC1 (48.2%) primarily separates control from ΔGwl samples, while PC2 (14.1%) distinguishes between developmental stages. Each point represents an individual biological replicate. Dashed lines indicating PC1 = 0 and PC2 = 0 axes. **(C)** PCA of significantly modulated phosphopeptides (t-test) of GVBD and GVBD+4h stages combined within each treatment group (control vs ΔGwl). Phosphopeptides significantly modulated (q < 0.05, two-group t-test with FDR correction) were used for log_₂_-transformed phosphorylation intensities. Each point represents a biological replicate, with control samples depicted in red (CT GVBD) and blue (CT GVBD+4H) and ΔGwl samples in green (ΔGwl GVBD) and grey (ΔGwl GVBD+4H). The percentage of variance explained by PC1 and PC2 is shown in parentheses. Dashed lines indicate PC1 = 0 and PC2 = 0. **(D)** Heat map illustrating the dynamic changes in differentially phosphorylated peptides between control (GVBD and 4 hours post GVBD) and Gwl-devoid oocytes (GVBD and 4 hours post GVBD).

We next checked the global proteome and phosphoproteome changes induced by Gwl depletion in meiotic oocytes. As depicted in the heatmap analysis of Figure 2D, the loss of Gwl promoted both dephosphorylation (upper heatmap part) and hyperphosphorylation (lower heatmap part) at a similar proportion (49,6 % vs 51,7 respectively) in numerous peptides of both GVBD and GVBD+4h groups. Although some differences were observed between these two stages (Figure 2D, red square), dephosphorylation/hyperphosphorylation was consistently widespread across all five biological replicates. Interestingly, we did not observe major changes in protein levels, with only 46 from a total of 4371 identified proteins displaying differential expression between control and ΔGwl oocytes. Interestingly, this short list included cyclin B, whose abundance was significantly higher in ΔGwl oocytes. These data suggest that phosphorylation modifications were not a consequence of changes in protein abundance and support a robust and reproducible effect of Gwl depletion in phosphoprotein pattern modification.

Our phosphoproteomic data identified 15,386 phosphopeptides from 5,300 proteins, from which 78.1% contained phospho-serine (pS), 20.8% phospho-threonine (pT), and 1.1% phospho-tyrosine (pY) (Figure 3A). From these phosphopeptides, we identified 12987 pS/pT unique sites with 20% displaying either increased or decreased phosphorylations upon Gwl depletion. Interestingly, we observed that 40,4% of the differential phosphorylations were performed in cyclin/CDK S/T-P consensus motifs indicating that 59,5% of pS/pT modifications were likely performed by other kinases/phosphatases indirectly modified by the newly established cyclin B/CDK1/PP2A-B55 equilibrium (Figure 3B). Our data additionally show a similar proportion of pTP and non-pTP differentially phosphorylated sites whereas pSP represented a half of the non-pSP motif suggesting that phosphorylation changes were preferentially performed in the TP motifs.

**Figure 3:**
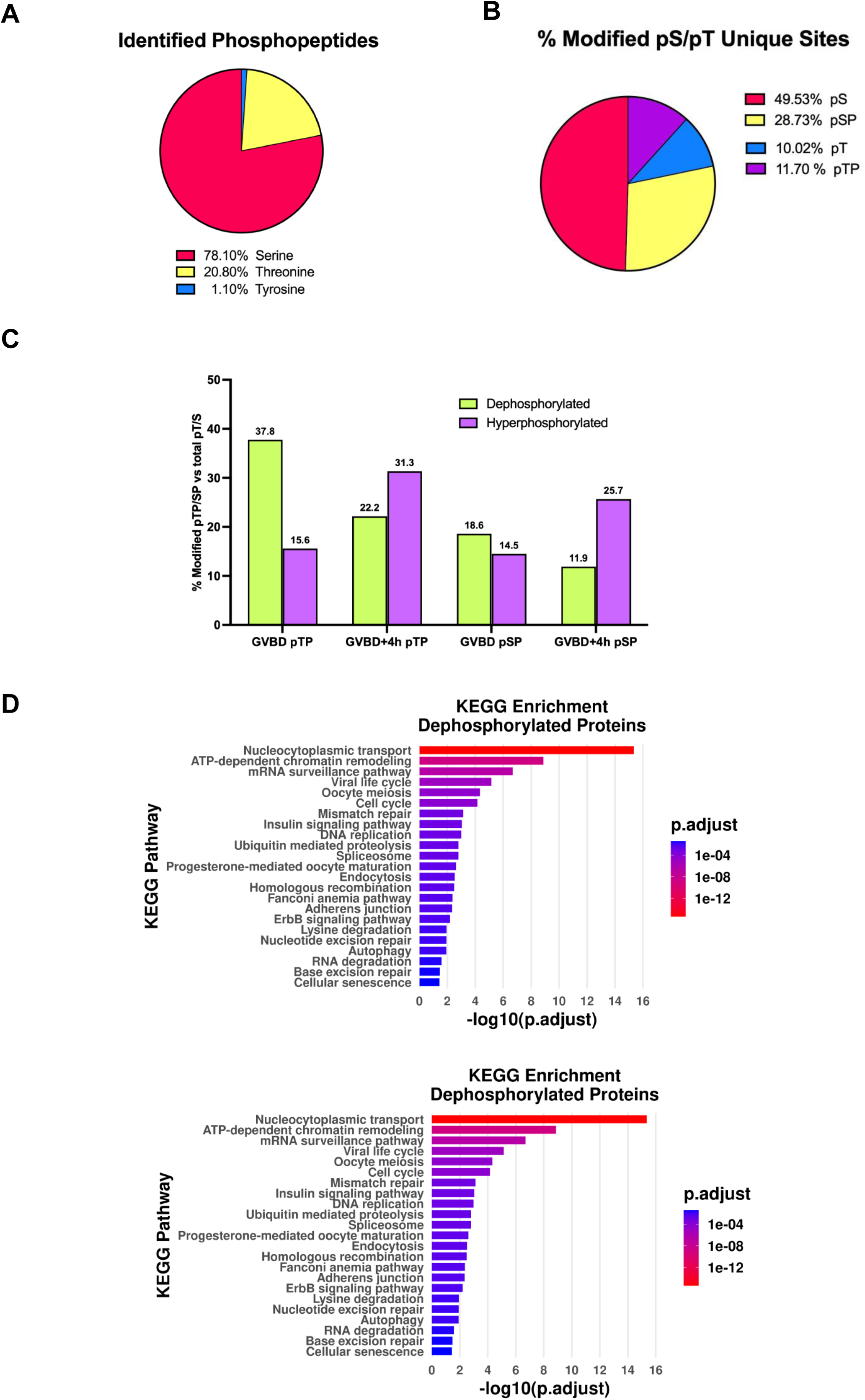
Analysis of differentially phosphorylated peptides in Gwl depleted oocytes. **(A)** Representation of the percentage of serine, threonine and tyrosine phosphosites identified in the label-free phosphoproteome. **(B)** Representation of the percentage of non-SP/TP and SP/TP unique sites differentially dephosphorylated (q<0.05) in control vs Gwl depleted oocytes of both GVBD and GVBD+4h conditions combined. **(C)** The percentage of unique SP and TP consensus sites at GVBD and GVBD+4H phosphorylated (violet) or dephosphorylated (green) upon Gwl depletion respect to total pS sites were represented. **(D)** KEGG pathway enrichment analysis of differentially dephosphorylated and phosphorylated proteins in control and ΔGwl oocytes in GVBD and GVBD+4h conditions combined.

Interestingly, at GVBD we observed a higher percentage of dephosphorylation than hyperphosphorylation in ΔGwl oocytes (Figure 3C). Considering that PP2A-B55 has a preference TP dephosphorylation^15,16^, these differences could reflect direct dephosphorylation of pTP sites by this phosphatase. Phosphorylation/dephosphorylation distribution was inversed in SP/TP cyclin B/CDK1 consensus motifs at GVBD+4h with a drastic increase in phosphorylation, notably in pSP consensus sites. Thus, Gwl depletion results in a marked dephosphorylation of pTP motifs at GVBD and hyperphosphorylation of pSP at GVBD+4h. Although alternative explanations cannot be ruled out, the observed dephosphorylation at GVBD could be attributable to a delay or failure of cyclin B/CDK1 to phosphorylate its substrates in the presence of hyperactive PP2A-B55 during early meiotic stages. Conversely, hyperphosphorylation at GVBD+4h may stem from an hyperactivation of cyclin B/CDK1 due to the accumulation of cyclin B and/or from the sequential activation of additional kinase cascades typically suppressed under control conditions.

Finally, we explored the potential biological functions of the differentially phosphorylated proteins using Kyoto Encyclopedia of Genes and Genomes (KEGG) pathway analysis. As expected, this analysis underscored an enrichment of hyperphosphorylated and dephosphorylated sites in proteins involved in progesterone-mediated oocyte meiosis, cell cycle and oocyte maturation confirming that Gwl plays an essential role in the control of meiotic maturation. However, we also identified other unexpected pathways such as nucleocytoplasmic transport, ATP-dependent chromatin remodelling, or mRNA surveillance, highlighting the involvement of Gwl in new unexpected cellular processes (Figure 3 D).

### Gwl depletion prevents c-Mos and Erp1 accumulation in maturing oocytes

To further investigate the role of Gwl in meiotic progression, we focused our interest in those dephosphorylated proteins known to be involved in the control of this cellular process. One major pathway controlling meiotic maturation is the c-Mos/MEK1/MAPK1 cascade. During meiotic progression, c-Mos is translated at GVBD promoting the activation of the MAP Kinase Kinase 1, MEK1, and the final phosphorylation of MAPK1^17^. Once activated, MAPK1 turns on Rsk1/2 (hereafter referred to as “Rsk”) which then phosphorylates and stabilizes Erp1, resulting in the subsequent inhibition of the APC/C and the establishment of the metaphase II arrest ^18^ (Figure 4A) . In our phosphoproteomic data, we identified T23 of MEK1 and Ser8, Ser26, T188, Y190 and T193 of MAPK1 as being dephosphorylated in Gwl-devoid Pg-treated oocytes (Figure 4B). Interestingly, T188 and Y190 belong to the TEY motif of MAPK1 and its double T188/Y190 phosphorylation promotes its activation^19,20^. Moreover, phospho-T193 is an autophosphorylation site acting as a feedback autoregulation to restrict MAPK1 activity^21^. We thus used specific phospho-antibodies against the TEY motif to confirm Y190 and T188 dephosphorylation upon Gwl depletion. As expected, phosphorylation of this motif was observed upon Pg addition in control oocytes from GVBD concomitantly with the activation of Plx1 (T210 phosphorylation) and of cyclin B/CDK1 (Y15 dephosphorylation). However, TEY phosphorylation was undetectable in Gwl depleted oocytes (Figure 4C) despite these oocytes activated cyclin B/CDK1, indicating that MAPK1 cannot be turned on in the absence of Gwl. To confirm that our findings resulted specifically from the loss of Gwl and not from unintended off-target effects, we performed a rescue experiment by re-expressing human Gwl (hGwl). Thus, we microinjected oocytes with HA-tagged TRIM21 mRNA, anti-Xenopus Gwl antibodies and purified human Gwl protein. Because human Gwl only shares 53% amino acid sequence identity with its Xenopus orthologue, it is barely recognized by anti-Xenopus Gwl antibodies and thus, cannot be targeted by TRIM21. As expected, under these conditions, MAPK1 was fully restored prior to GVBD in ΔGwl+hGwl oocytes supporting the specificity of our phenotype (Figure 4D).

**Figure 4:**
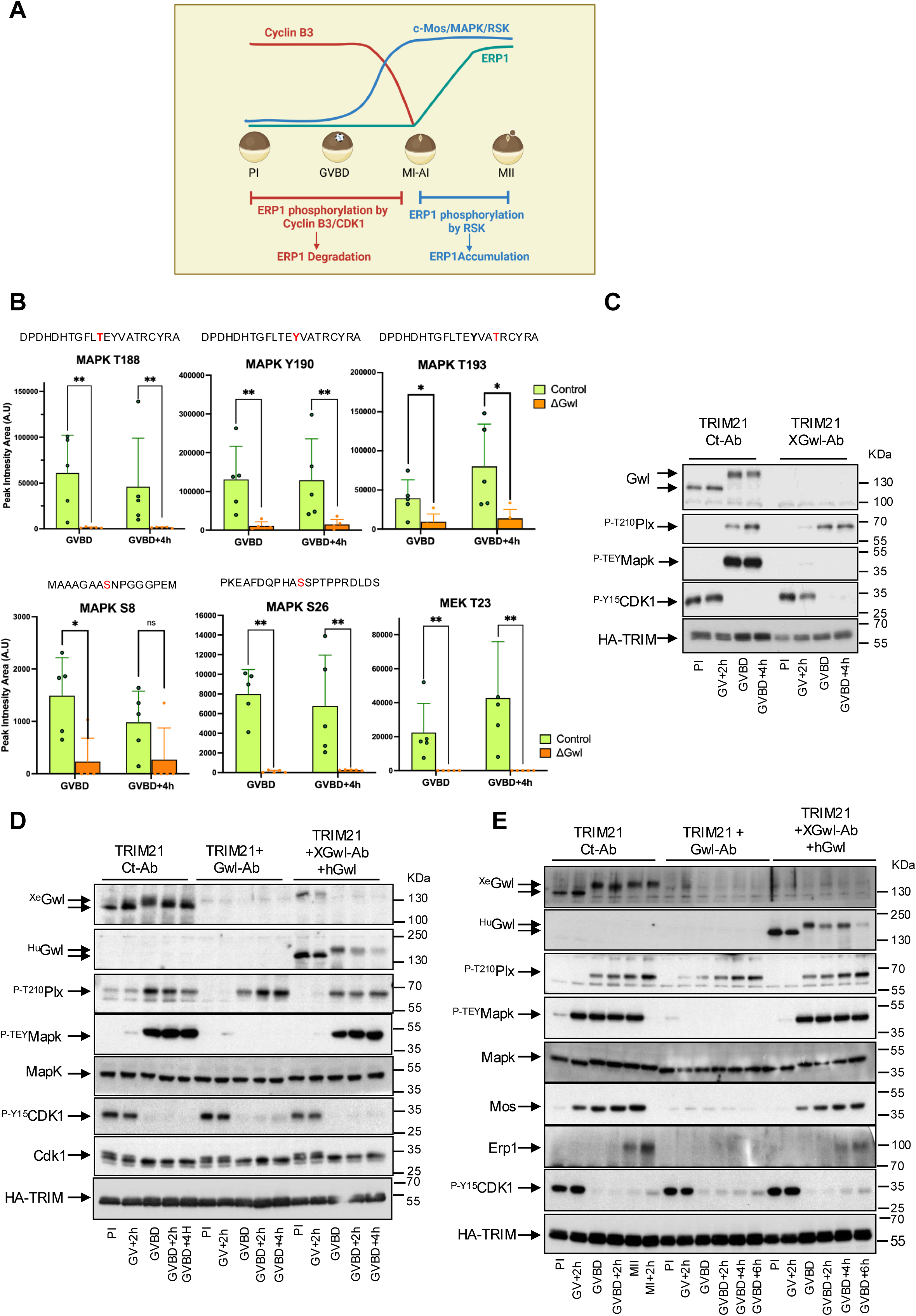
Gwl depletion prevents c-Mos and Erp1 accumulation in maturing oocytes. **(A)** Scheme illustrating the mechanisms controlling Erp1 stability during meiotic maturation. **(B)** Pick intensity area of the indicated phosphosites on MAPK and in MEK control and ΔGwl oocytes obtained by mass spectrometry in GVBD and GVBD+4h conditions. The position of the indicated phospho-sites in the TEY motif is indicated in red. Data are expressed as mean ± Standard Deviation (SD) from five independent replicates. Statistical analyses using two-sided non-parametric Mann-Whitney U test were performed. *p<0.05; **p<0.008; ****p<0.0001. n= 5. **(C)** Prophase I arrested oocytes were injected with HA-TRIM21 mRNA and control or Gwl antibodies. Sixteen hours later, oocytes were treated with Pg to induce meiotic maturation and the levels and phosphorylation of the indicated proteins analysed by western blot at the specified time points. **(D)** Prophase I oocytes were injected with HA-TRIM21 mRNA and control or Gwl antibodies. Sixteen hours later half of the oocytes devoid of Gwl were injected with the human Gwl protein (hGwl). One hour later oocytes were treated with Pg and immunoblotted at indicated time points to detect Xenopus (^Xe^Gwl) and Human Gwl (^Hu^Gwl), MAPK, CDK1 and TRIM21 protein levels and the phosphorylation of Plx1 on T210, MAPK on the TEY motif and of CDK1 on Y15. **(E)** Similar to (D) but c-Mos and Erp1 proteins were additionally examined. Data representative of at least three different experiments.

Next, we sought to determine whether the dephosphorylation of MAPK1 could result from the inactivation of its upstream kinases, MEK1 and c-Mos. We observed a dephosphorylation of T23 of MEK1 upon Gwl depletion (Figure 4B), however, the potential impact of the phosphorylation of this site on MEK1 activity is not known. Thus, we focused in the upstream kinase c-Mos. We repeated our Trim-Away experiment and we evaluated by western blot the levels and phosphorylation of the different meiotic proteins including MAPK1, c-Mos and Erp1. Interestingly, Gwl-depleted oocytes exhibit inactive MAPK1 and were devoid of c-Mos and Erp1 throughout the experiment, while control oocytes accumulated c-Mos from 2h upon Pg treatment and Erp1 upon GVBD (Figure 4E). Importantly, this effect is specific to Gwl depletion since the microinjection of hGwl restored c-Mos expression, reactivated MAPK, and stabilized Erp1 (Figure 4E). These data suggest that Gwl depletion in maturing oocytes inhibits c-Mos translation and/or stabilization and subsequent MEK1, MAPK1 and Rsk activation preventing the phosphorylation and accumulation of Erp1. If this is the case, the overexpression of c-Mos in Gwl depleted oocytes should restore MEK1/MAPK1/Rsk cascade activation and the final phosphorylation and stabilization of Erp1. However, surprisingly, the co-injection of c-Mos protein with TRIM21 and Xenopus-Gwl antibodies rescued MAPK1 phosphorylation but was unable to restore Erp1 protein levels (Figure 5A).

**Figure 5:**
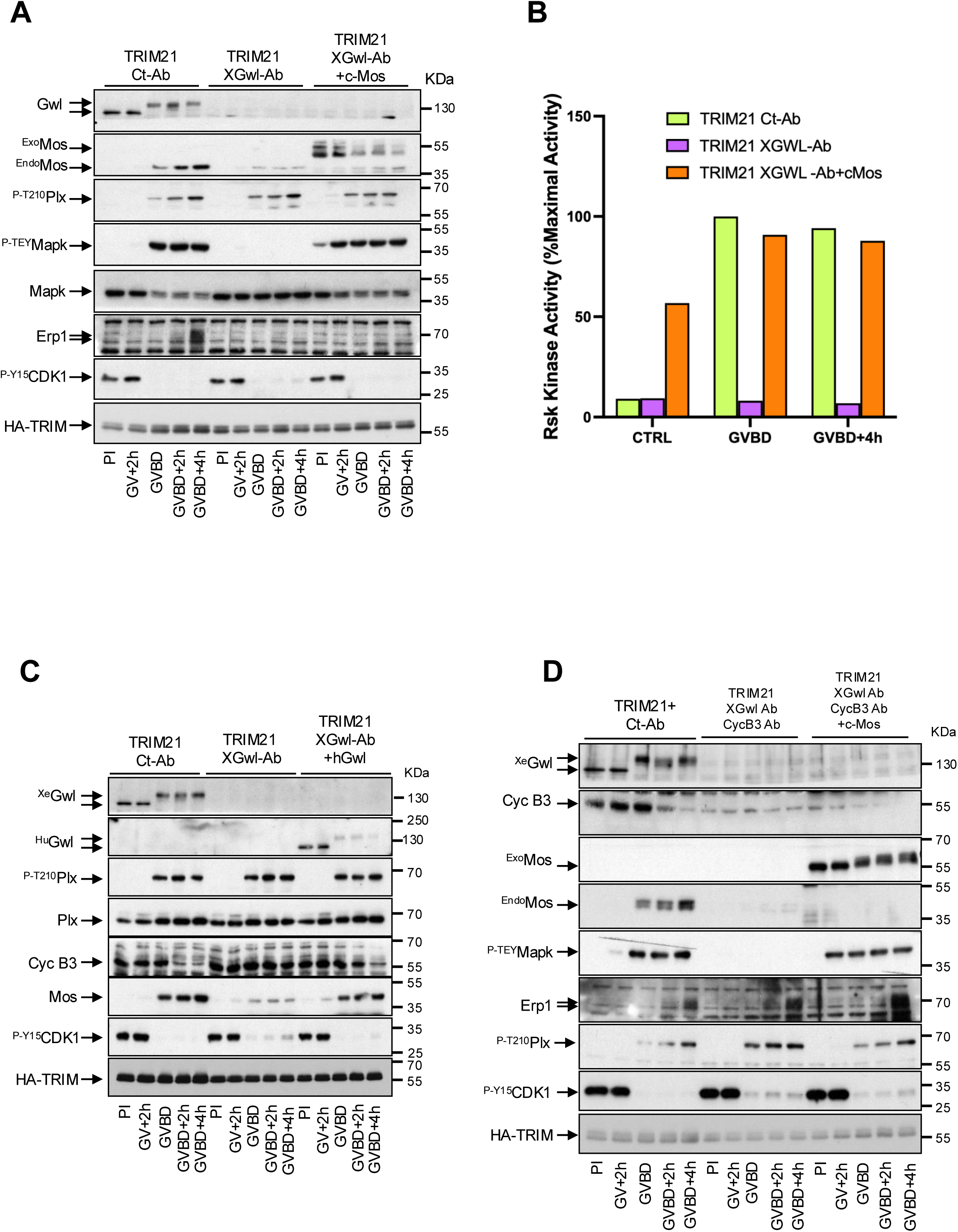
Xenopus oocytes lacking Gwl fail to accumulate c-Mos and Erp1 proteins and to degrade cyclin B3. **(A)** Prophase I oocytes were injected with HA-TRIM21 mRNA and control or XGwl Abs. Sixteen hours later, half of Gwl-devoid oocytes were injected with Xenopus c-Mos protein and subsequently treated with Pg. Oocytes were then recovered at the indicated time-points and immunoblotted to detect Gwl, endogenous and overexpressed tagged c-Mos, MAPK, Erp1 and TRIM21 protein levels as well as regulatory phosphorylation from Plx1 on T210, Mapk on the TEY motif and Cdk1 on Y15. **(B)** P90Rsk kinase activity was measured in vitro using the Kemptide peptide as a substrate in control, ΔGwl and ΔGwl+cMos oocytes and represented as the percentage of maximal activity. **(C)** Prophase I oocytes were injected with HA-TRIM21 mRNA and Control or XGwl antibodies. Sixteen hours later, half of the oocytes lacking the Gwl protein were injected with human Gwl protein and subsequently treated with Pg and analysed by Western blot at indicated time points to determine the phosphorylation and the levels of the specified proteins. **(D)** Xenopus prophase I arrested oocytes were injected with HA-TRIM21 mRNA and control or both XGwl and cyclin B3 antibodies. 16h later, half of the oocytes lacking Gwl and Cyclin B3 proteins were injected with Xenopus c-Mos protein. Upon treatment with Pg a sample of these oocytes were removed at the indicated time-points and immunoblotted to detect the levels and the phosphorylation of the specified proteins. Data representative of at least two different experiments.

The absence of Erp1 upon c-Mos co-injection could be due to the incapacity of the injected c-Mos protein to provide enough MAPK1 activity to turn on Rsk and to trigger Erp1 phosphorylation and accumulation or to the dysregulation in Gwl depleted oocytes of an additional pathway participating to Erp1 stability. To address these hypotheses, we evaluated Rsk activity under three experimental conditions: control, ΔGwl and ΔGwl+c-Mos. Our results clearly indicate that exogenous c-Mos effectively activates Rsk, however, this activation is unable to stabilize Erp1 in the absence of Gwl (Figure 5B). Our results clearly suggest that Gwl might regulate another pathway leading to Erp1 stabilization.

### Depletion of Greatwall from Xenopus oocytes impairs cyclin B3 degradation

Recent studies showed that cyclin B3-CDK1 kinase phosphorylates Erp1 from prophase I to anaphase I promoting its degradation by the proteasome and preventing the arrest of oocytes during metaphase I^4^. At anaphase I, this phosphorylation-dependent destabilization mechanism of Erp1 is halted by the degradation of cyclin B3. Thus, Erp1 stability is controlled by both Rsk- and cyclin B3-CDK1-dependent phosphorylation (Figure 4A). Data above demonstrate that Rsk activation is not sufficient to stabilize Erp1. Hence, we next checked whether Gwl depletion could perturb cyclin B3 stability during oocyte maturation. As expected, cyclin B3 is degraded at anaphase I in control oocytes, once MAPK1, cyclin B/CDK1 and Plx-1 are activated. However, this protein was stabilized throughout meiosis in Gwl depleted oocytes (Figure 5C). This effect appears to be specific since the addition of hGwl restores cyclin B3 degradation. These findings suggest that, besides the incapacity to synthesize c-Mos, the absence of Gwl in the oocytes prevents cyclin B3 proteolysis, a double effect that will prevent Erp1 stabilization and boost its degradation. To explore whether cyclin B3 degradation also contributes to the absence of Erp1 in ΔGwl oocytes, we performed a double Gwl/cyclin B3 Trim-Away experiment. We first confirmed that both proteins were effectively degraded in this experiment and we subsequently measured by western blot the levels and the phosphorylation of the meiotic proteins (Figure 5D). Interestingly, unlike c-Mos expression, the loss of cyclin B3 led to the accumulation of Erp1 upon GVBD in the absence of Gwl, an effect that was significantly enhanced when we supplemented c-Mos in our double Gwl/cyclin B3 Trim-Away assay. Importantly, in the absence of Gwl, c-Mos expression and Rsk activation appears not to be essential for Erp1 stabilization upon cyclin B3 degradation (Figure 5D). These data demonstrate that Gwl coordinates both, cyclin B3/CDK1 and c-Mos/MEK1/MAK1/Rsk cascade by modulating PP2A-B55 activity to confer the correct temporal pattern of Erp1 stabilization and MII arrest.

### Cyclin B3 degradation is mediated by the APC/C and conserved destruction motifs in Xenopus mitotic egg extracts

In order to further investigate the mechanisms by which Gwl depletion stabilizes cyclin B3, we next examined how cyclin B3 was degraded in mitotic Xenopus egg extracts. In human cells and mouse oocytes, cyclin B3 degradation takes place concomitantly with the activation of the APC/C, and the mutation of a putative Dbox motif (RXXFXXXXN) present in its amino acid sequence, induces its stabilization^22,23^. This strongly suggests that the APC/C complex may be the E3 ubiquitin ligase responsible for targeting cyclin B3 degradation. Interestingly, this putative Dbox sequence is also conserved in Xenopus cyclin B3 (Figure 6A). We thus, checked whether the mutation of this Dbox could also stabilize the protein in mitotic Xenopus egg extracts. To this, we evaluated the stability of either a wild-type or a Dbox arginine-to-alanine (R/A) mutant form of a [^35^S]-methionine-radiolabelled cyclin B3 in Xenopus egg extracts in which we induced mitotic entry by the addition of the active K72M mutant form of human Gwl (hK72MGwl) (Figure 6B). As expected, we observed that wild-type cyclin B3 was kept unchanged in interphase extracts whereas it was degraded 20 min upon hK72MGwl addition concomitantly with the degradation of endogenous cyclin B2. However, the R/A cyclin B3 mutant remained stable. Interestingly, the APC/C subunit APC3/Cdc27 displayed a normal phosphorylation and endogenous cyclin B2 degradation was not perturbed under these conditions, indicating that R/A mutant did not interfere with APC/C activity or with its capacity to degrade other cyclins. To further assess the role of the APC/C in cyclin B3 degradation, we next measured the stability of wild type cyclin B3 upon depletion of the APC/C subunit APC3/Cdc27 in these extracts. Confirming the role of the APC/C in cyclin B3 proteolysis, we observed its complete stabilization under these conditions (Figure 6C).

**Figure 6.**
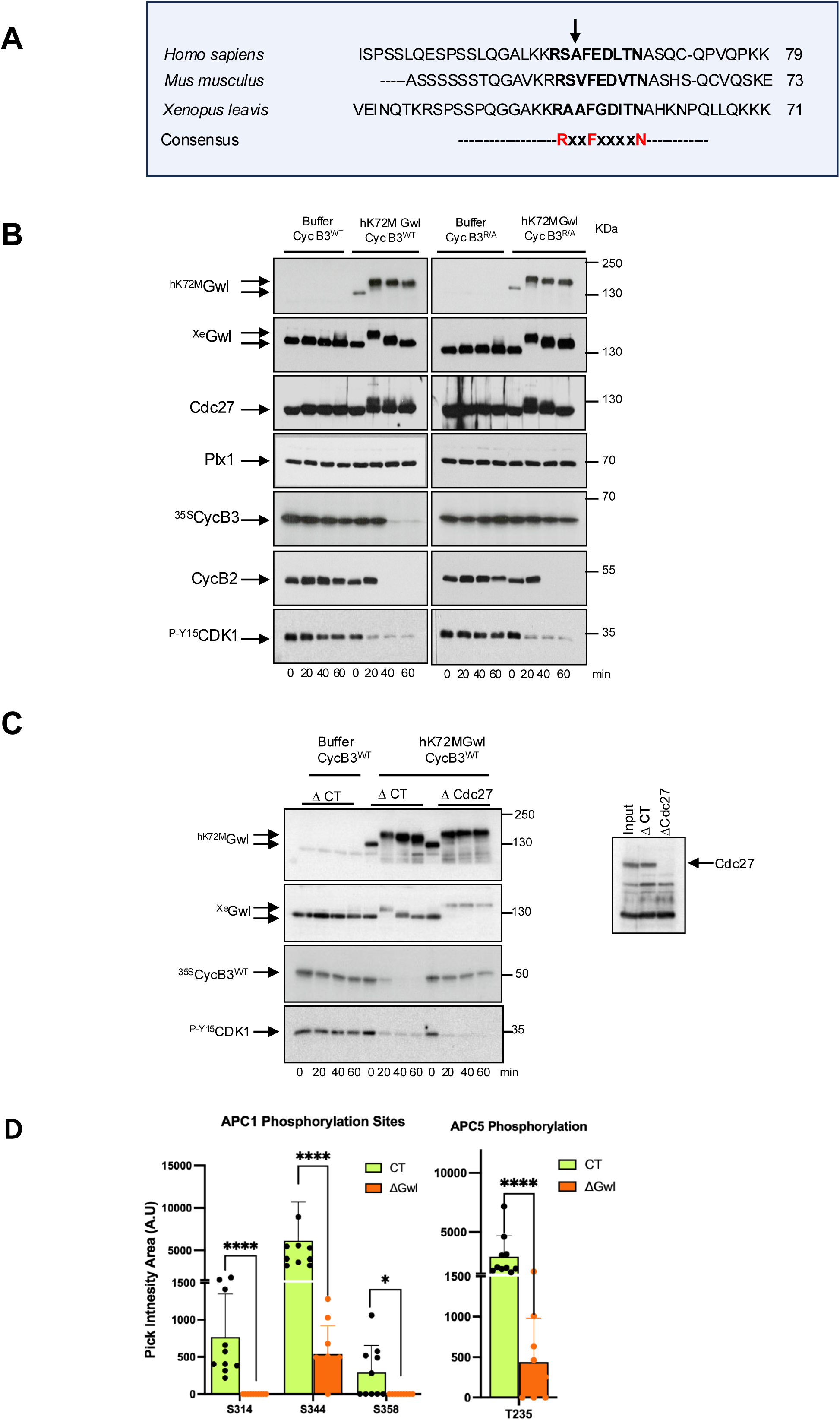
Cyclin B3 degradation is mediated by the APC/C. **(A)** Sequence alignment of the putative Dbox motifs of Cyclin B3 across different species. This motif is conserved in the Xenopus orthologue. **(B)** Interphase Xenopus egg extracts were supplemented with wildtype (WT) or mutated (R51A) in vitro translated ^35^S-Cyclin B3 protein in the presence or the absence of human Gwl K72M hyperactive kinase (hK72M). Samples were taken at indicated time-points and immunoblotted to examine human (^hK72M^Gwl) and Xe Gwl (^Xe^Gwl), Cdc27, Plx1, Cyclin B2 and phosphorylation of CDK1 on Y15. ^35S^Cyclin B3 was revealed by autoradiography. **(C)** Interphase Xenopus egg extracts were depleted of the APC3-Cdc27 protein and supplemented with in vitro translated ^35^S-Cyclin B3 protein. Samples were then used for immunoblotting. Cyclin B3 levels were revealed by autoradiography. Complete depletion of Cdc27 in these extracts was confirmed by western blot. Data representative for at least three different experiments. **(D)** Pick intensity area of the indicated phosphosites on the APC1 and APC5 subunits in control and ΔGwl oocytes obtained by mass spectrometry in GVBD and GVBD+4h combined conditions. Data are expressed as mean ± Standard Deviation (SD) from five independent replicates. Statistical analyses using two-sided non-parametric Mann-Whitney U test were performed. *p<0.032; ****p<0.0001.

### Greatwall modulates APC/C activation and orchestrates the temporal pattern of protein degradation also during meiosis in Xenopus oocytes

Data above demonstrates that in Xenopus egg extracts, cyclin B3 is degraded by the APC/C during mitotic division. The activity of this E3 ligase is controlled by the phosphorylation of different subunits of this complex including APC1, APC5 and APC3/Cdc27^24–26^. We thus hypothesized that the stabilization of cyclin B3 observed in Gwl-depleted maturing oocytes could result from a dephosphorylation and inactivation of the APC/C. Additionally, as supported by our proteomic findings, this inactivation would be also associated with the accumulation not only of cyclin B3, but also of cyclin B1 and B2 resulting in an increase of total cyclin/CDK activity. According to this hypothesis, our phosphoproteomic data highlighted a drastic dephosphorylation of APC1 at residues S314, S344 and S358, three essential phosphorylation sites controlling APC/C activity^26,27^, as well as an additional dephosphorylation of APC5 on T235 (Figure 6D). To confirm these data, we examined APC/C phosphorylation by western blot analysis in control and Gwl-devoid maturing oocytes. As expected, oocytes lacking Gwl displayed a dephosphorylation of APC3/Cdc27 and of APC1 in S358 and S314-318 sites as well as an accumulation of cyclin B1, B2 and B3 from GVBD (Figure 7A). Conversely, in control oocytes Cdc27/APC3 and APC1 were phosphorylated and cyclin B1, B2 and B3 were degraded from GVBD+2h. Because of the accumulation of cyclin B1, cyclin B2 and B3 in Gwl depleted oocytes, we also observed a significant rise of cyclin B/CDK activity upon GVBD (Figure 7B), however, this increase was not sufficient to phosphorylate APC/C subunits probably due to the hyperactivation of its counterbalancing phosphatase PP2A-B55. Finally, we checked whether these modified parameters were restored when hGwl was microinjected in Gwl Trim-Away-treated oocytes. Indeed, the ectopic addition of human Gwl in these extracts fully restored the control pattern, namely, the phosphorylation of the different APC/C subunits from GBVD, the partial degradation of cyclin B1 and B2 at GVBD, its re-accumulation at MII and the loss of cyclin B3 from GBVD+2h (Figure 7A). Together, these data strongly support, for the first time, the role of Gwl in conferring correct activation of the APC/C and highlights its critical function in orchestrating the temporal pattern of protein degradation during meiotic progression.

**Figure 7.**
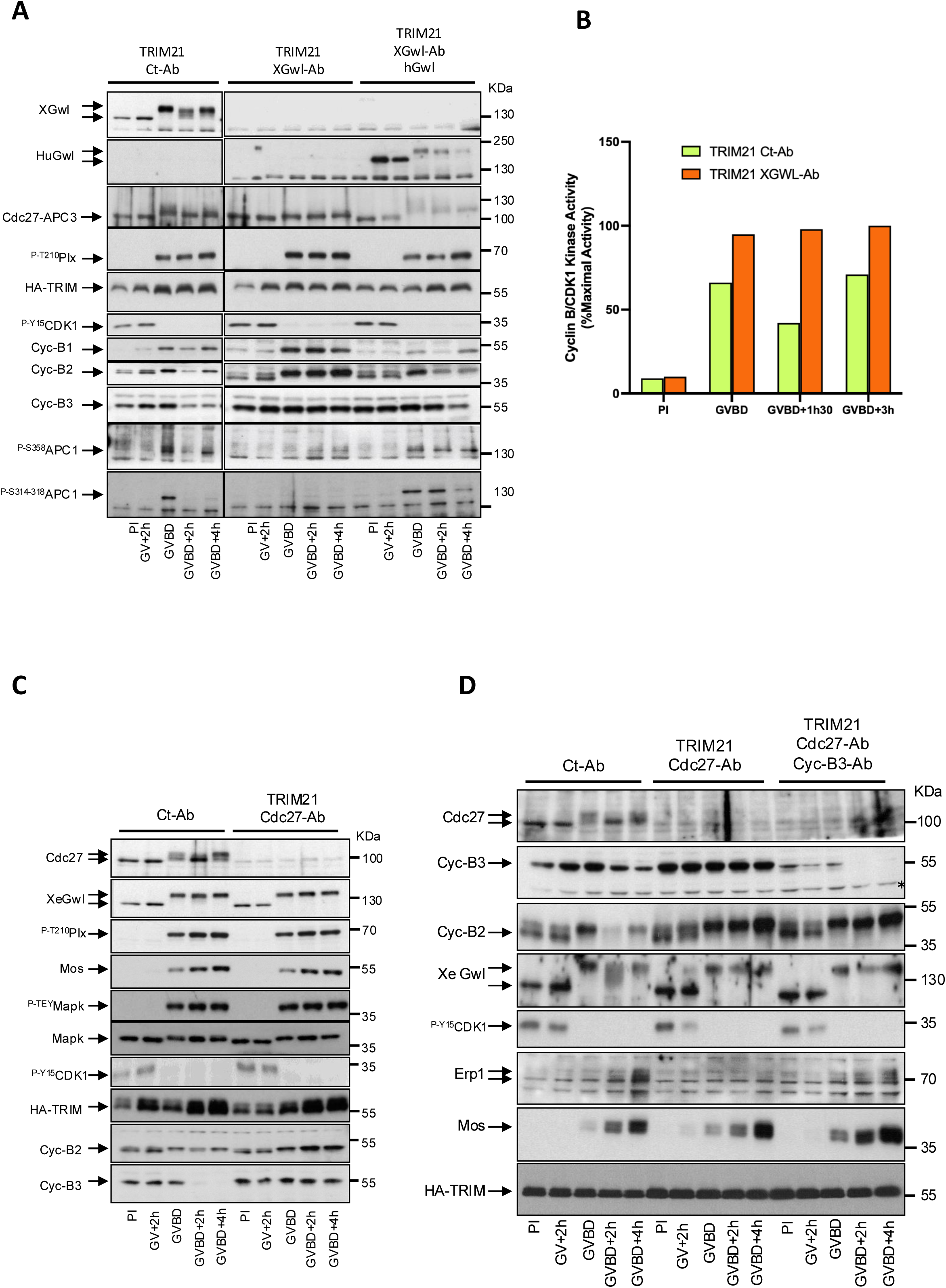
Inactive APC/C resulting from Gwl loss prevents Erp1 accumulation and the establishment of the metaphase II arrest by impairing cyclin B3 degradation. **(A)** The levels and the phosphorylation of the indicated proteins of maturing oocytes injected with control and XGwl antibodies and supplemented or not with the human Gwl protein were measured by western blot. Data representative of two different experiments. **(B)** Cyclin B/CDK1 kinase activity was measured in vitro in CDK1 immunoprecipitates from control or XGwl antibodies injected oocytes at the indicated time-points using H1 protein as a substrate. Values are represented as the percentage of maximal activity. **(C)** Prophase I arrested oocytes were injected with TRIM21 mRNA and control or anti-Cdc27 antibodies and treated with Pg. The levels and phosphorylation of the indicated proteins were analysed. **(D)** As for (C) except that a half of the anti-Cdc27 injected antibodies were also injected with anti-Cyclin B3 antibodies. Data representative of at least three different experiments.

### Inactive APC/C resulting from Gwl loss impairs cyclin B3 degradation and prevents Erp1 accumulation

Data above demonstrate that, during meiotic progression, the depletion of Gwl in oocytes does not allow the accumulation of Erp1 due to the absence of c-Mos and the stabilization of cyclin B3. They also demonstrate that the APC/C is not correctly activated in Gwl-devoid oocytes. Finally, we provide data showing that, as for cyclin B1 and B2, cyclin B3 is degraded via an APC/C-dependent ubiquitination. However, hitherto, we lacked direct evidence proving a causal relationship between inactive APC/C and the absence of Erp1 in these oocytes. Moreover, we have not data establishing whether inactive APC/C could modulate Erp1 levels directly, by promoting the accumulation of cyclin B3, or indirectly, by preventing c-Mos synthesis and MAPK1 activation. To obtain a definitive proof, we first inactivated APC/C in oocytes by APC3/Cdc27 Trim-Away and we evaluated the activation of c-Mos-MAPK1 during meiosis progression. Immunoblot analysis confirmed that c-Mos was correctly expressed and MAPK1 was fully phosphorylated in these oocytes, in which, as expected, cyclin B2 and B3 were accumulated attesting for the inactivation of the APC/C upon APC3/Cdc27 depletion (Figure 7C). These data prove that the APC/C does not modulate the c-Mos-MAPK1 pathway. Finally, we assessed the effect of the concomitant inactivation of the APC/C and depletion of cyclin B3 in maturing oocytes by performing a double APC3/Cdc27-cyclin B3 Trim-Away experiment. Importantly, as expected, the inactivation of the APC/C promoted cyclin B2 accumulation and the lack of Erp1, whereas the concomitant depletion of both the APC3 and cyclin B3 fully restored Erp1 accumulation in these oocytes (Figure 7D) indicating that perturbed APC/C activity stemming from Gwl loss prevents Erp1 stabilization by blocking cyclin B3 degradation. Thus, Gwl controls Erp1 accumulation by two independent pathways: (a) the modulation of c-Mos accumulation, and (b) the control of the APC/C via the degradation of cyclin B3.

## DISCUSSION

Our study identified new critical roles of Gwl and of cyclin B/CDK/PP2A-B55 balance in the control of meiotic progression in Xenopus oocytes. We used the Trim-Away approach to eliminate this kinase from prophase I-arrested Xenopus oocytes. This technique that, unlike morpholinos, directly induces an extremely efficient and specific degradation of the targeted protein, allowed us for the first time to fully eliminate Gwl kinase from prophase I-arrested Xenopus oocytes (note that Gwl is already present in stage VI oocytes). Unexpectedly, we first find that Gwl activity was not required to activate cyclin B/CDK1 in Pg-mediated meiotic resumption. This result contrasts with previously data showing that Gwl depletion by immunoprecipitation from G2 Xenopus egg extracts blocks mitotic entry by preventing Cdc25 phosphorylation and cyclin B/CDK1 activation^9^.

While the underlying cause of these differences remains unclear, we propose that the different balance between inhibitory Myt1/Wee1 kinases and activatory Cdc25 phosphatase in prophase I oocytes (expressing Myt1 but not Wee1) and G2 Xenopus egg extracts (expressing both Myt1 and Wee1) may play a significant role.

Indeed, PP2A-B55 phosphatase, prevents cyclin B/CDK1 activation by dephosphorylating Myt1, Wee1 and Cdc25 triggering the activation of the formers and the inhibition of the latter. If Myt1/Wee1 activity is higher than the one of Cdc25, cyclin B/CDK1 cannot be activated. Conversely, if Cdc25 activity surpasses Myt1/Wee1 inhibitory activity, cyclin B/CDK1 will be activated. In this line, as reported above, Wee1 is not expressed in prophase I-arrested oocytes and consequently CDK1 inhibitory activity may be lower in these oocytes^14^. Consistent with this notion, here we demonstrate that the expression of Wee1 in prophase I-arrested oocytes devoid of Gwl blocks GVBD. Moreover, we observe that the expression of Myt1 at similar levels does not prevent meiotic resumption indicating that Myt1 has a lower cyclin B/CDK1 inhibitory activity than Wee1. Thus, down-regulation of Wee1 translation before GVBD might be a key mechanism for the oocyte to permit prophase I-release by Pg.

Although cyclin B/CDK1 is activated upon Pg addition in Gwl-depleted prophase I arrested oocytes, we observed major protein phosphorylation changes during meiotic progression. Accordingly, our data showed that protein phosphorylation modifications, (hyperphosphorylation or dephosphorylation) were present in 20% of the phosphoproteome. Moreover, we could establish that 40% of modified phosphosites corresponded to S-T/P cyclin B/CDK1 consensus motifs known to be directly targeted by this kinase and by the PP2A-B55 phosphatase. As expected by the preference of PP2A-B55 for TP phosphorylated sites, we identified a higher percentage of dephosphorylated TP compared to SP motifs, whereas hyperphosphorylation was increased in SP sites. Interestingly, a higher percentage of dephosphorylation than hyperphosphorylation in both SP and TP motifs was present at GVBD, whereas this situation was inversed four hours later. These differences highlight the significant effect of the hyperactivation of PP2A-B55 stemming from Gwl depletion in meiotic progression. Considering our above data, we propose that the rapid hyperactivation of PP2A-B55 upon Gwl loss would delay phosphorylation of meiotic substrates till cyclin B protein levels will provide sufficient cyclin B/CDK1 threshold activity able to overpass phosphatase activity. This delay would explain protein dephosphorylation at GVBD. However, as meiosis progresses, and due to the incapacity of the APC/C to be phosphorylated and activated in the absence of Gwl, cyclin B1, B2 and B3 accumulates increasing total cyclin B/CDK1 activity and promoting hyperphosphorylation in late maturing oocytes (Figure S2 A). Moreover, besides the direct phosphorylation/dephosphorylation of meiotic substrates by cyclin B/CDK1 and PP2A-B55, the imbalance between these two enzymes might also promote indirect effects by activating some cascades that could participate to late protein hyperphosphorylation but, also, by inhibiting others that could be essential for the coordination of the different meiotic events. Accordingly, our data demonstrate that, besides abnormal high cyclin B/CDK1 activity in Gwl depleted oocytes, c-Mos is not accumulated and MAPK/Rsk pathway remains inactive in maturing oocytes. This dysregulation, together with an increase of cyclin B3 amount, prevented the stabilization of the inhibitor of APC/C, Erp1. Indeed, both c-Mos/MEK1/MAPK1/Rsk pathway and the cyclin B3/CDK1 kinase are essential to provide the correct temporal pattern of Erp1 protein expression during meiosis. Erp1 is dually phosphorylated by cyclin B3/CDK1 and by Rsk with the first one inducing its degradation until anaphase I and the second one promoting its stabilization from this stage of meiosis until fertilization^4,18,23^ (Figure 4A). Cyclin B3 is already present in prophase I-arrested oocytes and degraded upon anaphase I, when c-Mos starts to be accumulated. Thus, Erp1 is actively degraded before anaphase I and is further accumulated by Rsk-dependent phosphorylation from this stage to provide enough APC/C inhibitory activity to block oocytes in metaphase II.

Although the exact mechanisms that prevent c-Mos accumulation during meiotic resumption in Gwl-devoid oocytes have not yet been identified, they are likely to be the result of PP2A-B55-dependent dephosphorylation. It is possible that hyperactivation of PP2A-B55 stemming from Gwl depletion directly dephosphorylates c-Mos leading to its destabilization and degradation. In this line, we showed in the past that CDK1-dependent phosphorylation of c-Mos on S3 protects this protein from ubiquitin-mediated degradation^28^. In this scenario, increased PP2A-B55 activity may remove this stabilizing mark, making c-Mos susceptible to proteasomal degradation. Alternatively, c-Mos synthesis might be impaired. c-Mos mRNA translation is tightly regulated by the Cytoplasmic Polyadenylation Element-Binding protein (CPEB) that is essential for polyadenylation and translation activation^29^. The absence of Gwl might indirectly affect the phosphorylation of this translational regulator preventing efficient c-Mos synthesis. Supporting this idea, our phospho-mass spectrometry data revealed a significant dephosphorylation of CPEB.

Our data identified for the first time a primary cascade responsible for cyclin B3 stabilization upon Gwl depletion in maturing oocytes. Together, our phosphoproteomic and biochemical data demonstrate that in Gwl-devoid oocytes, APC1, APC5 and APC3/Cdc27 remain dephosphorylated during meiotic progression preventing the activation of this E3 ligase and impairing cyclin B1, B2 and B3 degradation at the MI-AI transition. As recently reported in mouse oocytes^30^, we identified cyclin B3 as a substrate of APC/C in Xenopus oocytes. More importantly, we also demonstrate that Gwl is essential for the activation of the APC/C and the degradation of cyclin B3, allowing the stabilisation of Erp1 crucial for the establishment of metaphase II arrest. Accordingly, when cyclin B3 was experimentally depleted in Gwl-deficient oocytes, Erp1 was restored, demonstrating that APC/C-mediated degradation of cyclin B3 plays a crucial role in maintaining Erp1 stability in these oocytes. These results reveal a novel feedback loop that is tightly regulated by Gwl/PP2A-B55. In this loop, the APC/C complex controls its own inhibitor, Erp1, by degrading its substrate, cyclin B3. Interestingly, although previous data showed that the stabilization of Erp1 at metaphase I upon cyclin B3 depletion requires c-Mos^4^, we demonstrate that this protein is not required in the absence of Gwl. In this regard, it has been shown that the presence of c-Mos during meiotic maturation promotes Rsk-dependent phosphorylation of Erp1 triggering its binding to PP2A and promoting the dephosphorylation of cyclin B/CDK1-dependent phosphosites that prime this protein for SCF-βTrcp-mediated ubiquitination and degradation^5^. Thus, in the absence of Gwl (e.g. in the presence of a hyperactive PP2A-B55), cyclin B/CDK1 phosphosites in Erp1 might be constantly dephosphorylated making Rsk activity dispensable for its stabilisation.

The drastic modification of protein phosphorylation and degradation stemming from Gwl loss makes difficult to define the meiotic stage in which Gwl-depleted oocytes are arrested. The inactivation of the APC and the hyperactivation of cyclin B/CDK1 in these oocytes, suggest that they would arrest in a metaphase I-like stage, however, because the absence of c-Mos, it is likely that additional key meiotic events regulated by the MAPK pathway are also missing in these oocytes. Finally, in the case that high PP2A-B55 activity permits these oocytes to proceed and finish meiosis I, they could not accumulate Erp1 and would be unable to arrest in metaphase II.

In summary, our data demonstrate that Gwl ensures the coordination of meiotic events by maintaining the proper balance between cyclin B/CDK1 kinase and PP2A-B55 phosphatase activities. This is achieved through the accurate temporal activation of APC/C and c-Mos accumulation, two events that are essential for meiosis I-II progression, restricting cyclin B3-dependent degradation of Erp1 prior to GVBD, and to stabilize Erp1 by the c-Mos/MEK1/MAPK1 cascade from this stage until fertilisation.

## METHODS

### EXPERIMENTAL MODEL AND SUBJECT DETAILS

#### *Xenopus laevis* Induction and Husbandry

Regulations for the use of *Xenopus laevis*, as outlined in the Animals Scientific Procedures Act (ASPA) and implemented by the Direction Generale de la Recherche et Innovation, Ministère de l’Enseignement Supérieur de la l’Innovation of France were followed.

Frogs were obtained from «Aquatic facilities: TEFOR Paris-Saclay», France and kept in a *Xenopus* research facility at the CRBM (Facility Center approved by the French Government. Approval n° C34-172-39).

Females were injected of 500 U Chorulon (Human Chorionic Gonadotrophin) and oocytes layed 18h later were used for experiments. Adult females were exclusively used to obtain eggs. All procedures were approved by the Direction Generale de la Recherche et Innovation, Ministère de l’Enseignement Supérieur de la l’Innovation of France (Approval n° APAFIS#40182-202301031124273v4).

#### Preparation of Interphase Xenopus Egg Extracts

Adult female Xenopus laevis were injected with 500 U of human chorionic gonadotropin (Chorulon) to induce ovulation. Eggs were collected approximately 18 h post-injection in XB buffer (100 mM KCl, 1 mM MgCl2, 0.1 mM CaCl2, 50 mM sucrose, 10 mM HEPES pH 7.8) at room temperature. Selected oocytes were dejellied using 2% cysteine solution (pH 7.8) for 5-7 minutes with gentle agitation. Metaphase II laid oocytes were treated with calcium ionophore A23187 (final concentration of 2 μg/ml) to induce meiotic exit and collected 35 minutes later. Extracts were prepared by centrifugation at 10,000 × g for 30 min at 4°C. The cytoplasmic fraction was collected, supplemented with protease inhibitors and centrifuged again at 10,000 × g for 30 min at 4°C. Supernatant was finally collected, aliquoted and frozen in liquid nitrogen for storage at -80°C.

#### Plasmids

Xenopus full length cyclin B3 cDNA was amplified from Xenopus ovary cDNAs, cloned blunt into EcoRV site of the pCS2 vector and subsequently subcloned either in the EcoRI-XbaI sites of pMalP2 or in the EcoRI-XhoI sites of pGEX4T2.

Full length human TRIM21 cDNA was cloned into the EcoRV cloning site of the HA-pCS2 vector. Full length Xenopus Myt1 and Wee1 cDNAs from Xenopus ovary cDNAs were cloned into the EcoRV cloning site of the HA-pCS2 vector.

Xenopus full length c-Mos was amplified from Xenopus ovary cDNAs and subcloned blunt into the EcoRV site of the pCS2 vector. This vector was subsequently used to subclone this sequence in EcoRI-NotI cloning site of pGEX4T1.

#### Site-Directed Mutagenesis

Point mutation of Xenopus cyclin B3 (R51A) was introduced by PCR in pCS2-Cyclin B3 plasmid using the Pfu ultra II fusion DNA polymerase (Agilent). Mutation was verified by DNA sequencing. Oligonucleotides used for site directed mutagenesis were purchased from Eurogentec.

#### mRNA preparation for Xenopus oocyte injection and in vitro translation

For expression of TRIM21, Wee1 or Myt1, corresponding pCS2 vectors were used to obtain the mRNAs using the mMESSAGE mMACHINE SP6 RNA polymerase Ultra Kit (Ambion) according to the manufacturer’s instructions.

Wild type cyclin B3 and R51A mutant mRNAs were transcribed from pCS2-Cyclin B3 (Wt and R51A) vectors with the mMessage mMachine SP6 RNA polymerase kit (Ambion) and translated in vitro using reticulocyte lysate system (Promega).

#### Antibody Production and Purification and Immunoblots

MBP or GST fusion cyclin B3 proteins were expressed in *E. Coli*. For the immunization protocol, GST-Cyclin B3 was purified from inclusion bodies, subjected to SDS–PAGE and electroeluted according to standard procedures. This protein was next dialysed against 50 mM NaCl, 50 mM NaHCO3 buffer and used to immunize rabbits. Soluble MBP-Cyclin B3 was purified in an amylose column and coupled to CNBr-sepharose beads for affinity purification of immune sera as previously described by Lorca and colleagues (Lorca et al., 1998).

GST-fused Xenopus c-Mos protein was expressed in *E. Coli* and purified using a gluthatione column. Purified c-Mos was then dialyzed against XB buffer, aliquoted and stored at -80°C. For western blot, 1 µl of interphase or maturing oocyte extracts was loaded on a polyacrylamide gel and transferred onto Immobilon-P membrane that was next blocked with 5% milk-TBST, or with 5% BSA-TBST when phospho-antibodies were used, and incubated overnight with primary antibodies in 2% milk-TBST. HRP conjugated secondary antibodies directed against rabbit (Cell Signaling) or mouse (Santa Cruz biotechnologies) were incubated for 30 min at RT in 2% milk-TBST.

#### TRIM-Away in Xenopus oocytes

Prophase I-arrested oocytes were harvested after surgery, washed in OR-2 buffer (82.5 mM NaCl, 2 mM KCl, 1 mM MgCl2, 5 mM HEPES, pH 7.2) and incubated with 1 mg/ml of collagenase. After dissociation, oocytes were selected, washed again in OR-2 buffer and injected with 20 nl of a mixture containing 10 ng of mRNA TRIM21 and 40 ng of control (anti-GST) or affinity purified (Xenopus Gwl, cyclin B3 or Cdc27) antibodies. For complementation studies, 20 ng of human wildtype Gwl and/or 40 ng of Xenopus c-Mos protein were subsequently injected. Upon injection, oocytes were incubated overnight at 19°C in NDS medium (96 mM NaCl, 2 mM KCl, 1.8 mM CaCl2, 1 mM MgCl2, 5 mM HEPES, 2.5 mM Pyruvate, 0.05 mM Gentamycine) and the day after, progesterone was added to a final concentration of 37 µM in Merriam buffer (10 mM HEPES, 0.82 mM MgSO4, 88 mM NaCl, 0.33 mM Ca(NO3)2, 1 mM KCl and 0.41 mM CaCl2, pH7.4). Oocytes in prophase I, 2 hours upon progesterone addition, at GVBD (assessed by pigment rearrangement in the animal pole) and 4 hours after GVBD were recovered. Collected oocytes were then homogenized in extraction buffer (80 mM β-glycerophosphate, 20 mM EGTA, 15 mM MgCl2 and 2 mM EDTA), centrifuged at 15,000 × g for 10 min at 4 °C and lysates used for Western blotting.

#### Immunoprecipitation

For immunoprecipitation, 2 μg of anti-CDK1 or anti-Rsk antibodies were incubated with 20 μl Dynabeads protein G (Invitrogen) for 30 min and, after washing with XB buffer, incubated again with 80 μl of the lysate corresponding to four oocytes for an additional period of 30 min. Beads were then washed to eliminate non-specific bound proteins and used for kinase activity assays. For immunodepletion, 10 μg of affinity-purified antibodies against APC3-Cdc27 or GST (Glutathione S-Transferase) were bound for 30 min to 20 μl Dynabeads protein G (Invitrogen), and then, after washing with XB buffer, added to 20 μl of Interphase egg extracts for 30 min at 20°C. Two consecutive immunoprecipitations were performed to completely remove endogenous Cdc27 protein. Then, beads were removed and the supernatant was incubated with radiolabelled cyclin B3 at 20°C. Samples were taken every 20 minutes up to 60 minutes, mixed 1:9 with 2x Laemmli buffer and heated 5 minutes at 94°C to be subsequent used for immunoblot analysis.

#### Kinase Activity Assays

CDK1 kinase activity was measured using histone 1 as a substrate. Shortly, four-oocyte CDK1 immunoprecipitates previously obtained and frozen, were thawed by the addition of 15 μl of histone buffer (20 mM Hepes pH 7.2, 5 mM MgCl2, 100 µM ATP and 1 mg/ml Histone 1), including 0.2μCi of [γ33P] ATP and incubated for 10 minutes at room temperature. Reactions were stopped by adding 2x Laemmli sample buffer followed by heat denaturation at 94°C for 5 minutes and analysed by SDS-PAGE and autoradiography.

For Rsk kinase activity measurements, Rsk1/2 immunoprecipitates from 4 oocytes were mixed with a buffer containing 1 mg/ml of dephosphorylated Kemtide peptide, 0.3 μM [γ33P] ATP, 2 mM MgCl2 and 50 mM Tris-HCL pH7.5 and incubated for 20 minutes at room temperature. Each reaction mixture was spotted onto P81 phosphocellulose paper squares (1 × 1 cm), dried for 1-2 minutes and washed 5 times (5 minutes each) with 75 mM phosphoric acid with gentle agitation to remove unincorporated [γ-³³P] ATP. A subsequent wash with acetone was performed and papers were allowed to dry completely. Dried papers were then placed in scintillation vials, scintillation liquid was added, and ³³P incorporation was measured using a scintillation counter.

#### Degradation Assay

For protein degradation assays, 2 μl of either ^35^S-labelled WT cyclin B3 or mutant (R51A) translated proteins, were incubated at room temperature with 20 μl of interphase extracts supplemented or not with Hu Gwl K72M protein.

#### Incubation and Meiotic Induction

Injected oocytes were incubated overnight at 19°C in NDS medium. Upon overnight incubation, oocytes were transferred to Merriam Buffer supplemented with progesterone (37 μM) to induce meiotic resumption.

#### Sample Preparation Prior to LC-MS/MS Analysis

Ten oocytes per condition were homogenized in extraction buffer, centrifuged at 15 000 x g for 10 min at 4°C and supernatants recovered. A six-time volume of cold acetone (−20°C) was then added to the samples containing about 250 µg of protein extracts. Vortexed tubes were incubated overnight at -20°C then centrifuged for 10 min at 12 900 x g at 4°C. Supernatant was removed, then the protein pellets were dissolved in 8 M Urea-25 mM NH4HCO3 buffer. Samples were then reduced with 10 mM TCEP-HCl and alkylated with 20 mM MMTS. After a 16-fold dilution in NH4HCO3, samples were then digested overnight at 37°C by a mixture of trypsin/Lys C (1/100 Enzyme/Substrate ratio). The digested peptides were desalted on Sep-Pak classic C18 cartridges. Cartridges were sequentially washed with 2 ml methanol, 2 ml of a 70% (v/v) aqueous ACN containing 1% (v/v) TFA and equilibrated with 2 ml of 1% (v/v) aqueous TFA. The digested peptides were acidified with 1% (v/v) aqueous TFA, applied to the column and washed with 2 x 1 ml of a 1% (v/v) aqueous FA. Peptides were eluted by applying 2 x 500 µL of a 70% (v/v) aqueous ACN containing 0.1% (v/v) FA. Eluates were vacuum-dried. Phosphopeptide enrichment was then performed according to manufacturer’s procedure starting from the lyophilized peptide samples. Eluted phosphopeptides and peptides from total proteome were loaded and desalted on evotips provided by Evosep (Odense, Denmark) according to manufacturer’s procedure before LC-MS/MS analysis. The flowthrough material resulting from the enrichment procedure was processed as well for total proteome analysis.

#### LC-MS/MS acquisition

Samples were analyzed on a timsTOF Pro 2 mass spectrometer (Bruker Daltonics, Bremen, Germany) coupled to an Evosep one system (Evosep, Odense, Denmark) operating with the 30SPD method developed by the manufacturer. Briefly, the method is based on a 44-min gradient and a total cycle time of 48 min with a C18 analytical column (0.15 x 150 mm, 1.9 µm beads, ref EV-1106) equilibrated at 40°C and operated at a flow rate of 500 nl/min. H2O/0.1 % FA was used as solvent A and ACN/ 0.1 % FA as solvent B. For the phosphopeptide analysis, the timsTOF Pro 2 was operated in DDA-PASEF mode (MeierF and Colleagues, 2015) with a cycle time of 1.3 seconds. Mass spectra for MS and MS/MS scans were recorded between 100 and 1700 *m/z*. The total proteome was also analyzed on the instrument using a DIA-PASEF method consisting of 12 pydiAID frames with 3 mass windows per frame, resulting in a cycle time of 0.975 seconds as described in the Bruker Application Note LCMS 218. Collisional energy was ramped stepwise as a function of ion mobility.

#### Data analysis

MS raw files for phosphopeptide analysis were processed using PEAKS Online 11 (build 1.9, Bioinformatics Solutions Inc.). Data were searched against the SwissProt+TrEMBL Xenopus Laevis entries (downloaded January 2024, 112365 entries). Parent mass tolerance was set to 20 ppm, with fragment mass tolerance to 0.05 Da. Specific tryptic cleavages were selected and a maximum of 2 missed cleavages were allowed. The following post-translational modifications were considered for identification: Phosphorylation (STY), Oxidation (M), Deamidation (NQ) and Acetylation (Protein N-term) as variable and Beta-methylthiolation (C) as fixed. Identifications were filtered based on a 1% FDR (False Discovery Rate) threshold at both peptide and protein group levels. Label free quantification was performed using the PEAKS Online 11 quantification module, allowing a mass tolerance of 5 ppm, a CCS error tolerance of 0.01 and a 0.25-min retention time shift tolerance for match between runs. PEAKS Studio software was also used to format sequences with the modified amino acid flanked by +/-10 amino acids.

DIA raw files for total proteome were processed using Spectronaut 18 (Biognosys, Switzerland). Data were searched against the same database. Specific tryptic cleavages were selected and a maximum of 2 missed cleavages were allowed. The following post-translational modifications were considered for identification: Acetyl (Protein N-term), Oxidation (M), as variable and MMTS (C) as fixed. The maximum number of variable modifications was set to 5. Identifications were filtered based on a 1% precursor and protein Qvalue cutoff threshold. The protein LFQ method was set to automatic and the quantity was set at the MS2 level with a cross-run normalization applied.

Multivariate statistics on protein or peptide measurements were performed using Qlucore Omics Explorer 3.9 (Qlucore AB, Lund, *SWEDEN*). A positive threshold value of 1 was specified to enable a log2 transformation of abundance data for normalization *i.e.* all abundance data values below the threshold will be replaced by 1 before transformation. The transformed data were finally used for statistical analysis *i.e.* evaluation of differentially present proteins or peptides between two groups using a Student’s bilateral t-test. Since the observed effects of Gwl kinase on phosphorylation are very close in the early and late conditions, we also statistically processed the samples considering only two Ctrl and Gwl groups of 10 replicates each (GVBD plus MII) and substracting the variance related to the timing parameter. A p-value better than 0.05 was finally used to filter differential candidates.

#### Phosphoproteomics data processing and filtering

Phosphoproteomic data were obtained from the file 2402007_ptmprofile_LFQ_processed doublons_annotations_Ttest CTRL vs Gwl-EF Timing-1.xlsx, which was processed in R (v4.3) using the packages {readxl}, {dplyr}, {purrr}, {stringr}, {magrittr}, {openxlsx}, {ggplot2}, {ggfortify}, {ggsci} and {ggrepel}. The data was imported with read_xlsx, and phosphorylation sites were annotated using columns for peptide sequences, protein IDs, gene names, amino acid type, and site position.

#### Phosphosite Filtering and Clustering

To group phosphorylation sites by protein, gene names were cleaned to remove isoform suffixes, and unique clusters were assigned to each protein (one cluster per unique gene). Sites were filtered to retain only unique phosphorylation sites per protein for each amino acid type: serine (S), threonine (T), and tyrosine (Y). This ensures that each phosphosite is only counted once per protein.

#### Analysis of Variance (ANOVA) and Multiple Testing Correction

For each retained peptide, an ANOVA was performed to assess differences in phosphorylation levels across the four experimental conditions. Resulting p-values were adjusted for multiple testing using the false discovery rate (FDR) method (Benjamini-Hochberg). Additionally, q-values were computed using the qvalue R package to estimate the proportion of false positives (Storey & Tibshirani, 2003).

#### Principal component analysis (PCA)

To explore the global variation in phosphorylation profiles, PCA was performed using the log2-transformed intensities of peptides significantly modulated across conditions (q < 0.05). Peptides were considered significantly modulated if they exhibited evidence of differential phosphorylation based on ANOVA followed by FDR correction. Statistical computing and graphics was performed using R software (https://www.R-project.org/).

The PCA was performed on a matrix where rows represent samples (biological replicates) and columns represent significantly modulated phosphopeptides. Principal components were calculated using the prcomp() function with centering and scaling applied to each peptide.

The PCA scores for each sample were visualized using ggplot2 with colour coding to indicate experimental condition. Sample labels were added using ggrepel to minimize overlap. Axes were annotated with the percentage of variance explained by each principal component.

#### Heat Map

Heat mat was performed using the ComplexHeatmap package (Gu and Collegues, 2016).

### QUANTIFICATION AND STATISTICAL ANALYSIS

Differences of phosphorylation in phosphorylation levels across the different conditions in label-free mass spectrometry analysis were performed using ANOVA followed FDR correction. When PCA was performed combining GVBD and GVBD+4H groups in control and ΔGwl conditions, a two-group t-test with FDR correction was used. Statistical differences in peak intensity area between GVBD and GVBD+4H of phosphopeptides described in Figure 4A and Figure 6D were determine by using a two-sided non-parametric Mann-Whitney U test.

### RESOURCE AVAILABILITY

#### Lead contact

Further information and requests for resources and reagents should be directed to and will be fulfilled by the lead contact

## Supporting information

Supplementary Informations

## ACKNOWLEDGMENTS

We would like to thank Marc Plays and Anthony Ollier for providing the Xenopus females, the ZEFIX animal facility and for immunising the rabbits and recovering the antibodies. APC1 phospho-antibodies were a generous gift from the laboratory of Hiro Yamano, University College of London, UK. TRIM21 cDNA was kindly provided by Sebastian Nisole, IRIM, Montpellier. The work in the laboratory was supported by grants from the Agence Nationale pour la Recherche (ANR, France ANR-20-CE13-0033), Université de Montpellier, ”Accelerateur Innovation” 2022 (NewObTrea) and the Fondation ARC pour la Recherche sur le Cancer (PJA3 TL ARCPJA2023060006673).

## AUTHOR CONTRIBUTIONS

S. R. Conceptualization; Resources; Data curation; Formal analysis; Validation; Investigation; Visualization; Methodology; Writing—review and editing. C.H.K. Formal analysis; Validation; Investigation; Methodology. C.B.C. Investigation, Data curation S.V. Investigation, Data curation. V.L. Formal analysis; Validation; Investigation; Methodology. G.C. Formal analysis; Validation; Investigation; Methodology. B.L. Formal analysis; Investigation; Methodology. A.C. Conceptualization; Resources; Formal analysis; Supervision; Funding acquisition; Validation; Visualization; Methodology; Writing—review and editing. T.L. Conceptualization; Resources, Data curation; Formal analysis; Supervision; Funding acquisition; Validation; Investigation; Visualization; Methodology; Project administration; Writing—review and editing.

## REFERENCES

1. Pei, Z., Deng, K., Xu, C., and Zhang, S. (2023). The molecular regulatory mechanisms of meiotic arrest and resumption in Oocyte development and maturation. Reprod Biol Endocrinol 21, 90. 10.1186/s12958-023-01143-0.

2. Ohe, M., Inoue, D., Kanemori, Y., and Sagata, N. (2007). Erp1/Emi2 is essential for the meiosis I to meiosis II transition in Xenopus oocytes. Dev Biol 303, 157–164. 10.1016/j.ydbio.2006.10.044.

3. Schunk, R., Halder, M., Schäfer, M., Johannes, E., Heim, A., Boland, A., and Mayer, T.U. (2025). A phosphate-binding pocket in cyclin B3 is essential for XErp1/Emi2 degradation in meiosis I. EMBO reports 26, 768–790. 10.1038/s44319-024-00347-8.

4. Bouftas, N., Schneider, L., Halder, M., Demmig, R., Baack, M., Cladière, D., Walter, M., Al Abdallah, H., Kleinhempel, C., Messaritaki, R., et al. (2022). Cyclin B3 implements timely vertebrate oocyte arrest for fertilization. Dev Cell 57, 2305–2320.e6. 10.1016/j.devcel.2022.09.005.

5. Wu, J.Q., Hansen, D.V., Guo, Y., Wang, M.Z., Tang, W., Freel, C.D., Tung, J.J., Jackson, P.K., and Kornbluth, S. (2007). Control of Emi2 activity and stability through Mos-mediated recruitment of PP2A. Proceedings of the National Academy of Sciences 104, 16564– 16569. 10.1073/pnas.0707537104.

6. Castro, A., and Lorca, T. (2018). Greatwall kinase at a glance. J Cell Sci 131, jcs222364. 10.1242/jcs.222364.

7. Lorca, T., Bernis, C., Vigneron, S., Burgess, A., Brioudes, E., Labbé, J.-C., and Castro, A. (2010). Constant regulation of both the MPF amplification loop and the Greatwall-PP2A pathway is required for metaphase II arrest and correct entry into the first embryonic cell cycle. J Cell Sci 123, 2281–2291. 10.1242/jcs.064527.

8. Zhao, Y., Haccard, O., Wang, R., Yu, J., Kuang, J., Jessus, C., and Goldberg, M.L. (2008). Roles of Greatwall kinase in the regulation of cdc25 phosphatase. Mol Biol Cell 19, 1317–1327. 10.1091/mbc.e07-11-1099.

9. Yu, J., Zhao, Y., Li, Z., Galas, S., and Goldberg, M.L. (2006). Greatwall kinase participates in the Cdc2 autoregulatory loop in Xenopus egg extracts. Mol Cell 22, 83–91. 10.1016/j.molcel.2006.02.022.

10. Mochida, S., Maslen, S.L., Skehel, M., and Hunt, T. (2010). Greatwall phosphorylates an inhibitor of protein phosphatase 2A that is essential for mitosis. Science 330, 1670–1673. 10.1126/science.1195689.

11. Gharbi-Ayachi, A., Labbé, J.-C., Burgess, A., Vigneron, S., Strub, J.-M., Brioudes, E., Van-Dorsselaer, A., Castro, A., and Lorca, T. (2010). The substrate of Greatwall kinase, Arpp19, controls mitosis by inhibiting protein phosphatase 2A. Science 330, 1673–1677. 10.1126/science.1197048.

12. Clift, D., So, C., McEwan, W.A., James, L.C., and Schuh, M. (2018). Acute and rapid degradation of endogenous proteins by Trim-Away. Nat Protoc 13, 2149–2175. 10.1038/s41596-018-0028-3.

13. Clift, D., McEwan, W.A., Labzin, L.I., Konieczny, V., Mogessie, B., James, L.C., and Schuh, M. (2017). A Method for the Acute and Rapid Degradation of Endogenous Proteins. Cell 171, 1692–1706.e18. 10.1016/j.cell.2017.10.033.

14. Nakajo, N., Yoshitome, S., Iwashita, J., Iida, M., Uto, K., Ueno, S., Okamoto, K., and Sagata, N. (2000). Absence of Wee1 ensures the meiotic cell cycle in Xenopus oocytes. Genes Dev 14, 328–338.

15. Kruse, T., Garvanska, D.H., Varga, J.K., Garland, W., McEwan, B.C., Hein, J.B., Weisser, M.B., Benavides-Puy, I., Chan, C.B., Sotelo-Parrilla, P., et al. (2024). Substrate recognition principles for the PP2A-B55 protein phosphatase. Science Advances 10, eadp5491. 10.1126/sciadv.adp5491.

16. Swartz, S.Z., Nguyen, H.T., McEwan, B.C., Adamo, M.E., Cheeseman, I.M., and Kettenbach, A.N. (2021). Selective dephosphorylation by PP2A-B55 directs the meiosis I-meiosis II transition in oocytes. eLife 10, e70588. 10.7554/eLife.70588.

17. Balmanno, K., Kidger, A.M., Byrne, D.P., Sale, M.J., Nassman, N., Eyers, P.A., and Cook, S.J. (2023). ERK1/2 inhibitors act as monovalent degraders inducing ubiquitylation and proteasome-dependent turnover of ERK2, but not ERK1. Biochem J 480, 587–605. 10.1042/BCJ20220598.

18. Inoue, D., Ohe, M., Kanemori, Y., Nobui, T., and Sagata, N. (2007). A direct link of the Mos-MAPK pathway to Erp1/Emi2 in meiotic arrest of Xenopus laevis eggs. Nature 446, 1100– 1104. 10.1038/nature05688.

19. Guadagno, T.M., and Ferrell, J.E. (1998). Requirement for MAPK activation for normal mitotic progression in Xenopus egg extracts. Science 282, 1312–1315. 10.1126/science.282.5392.1312.

20. Canagarajah, B.J., Khokhlatchev, A., Cobb, M.H., and Goldsmith, E.J. (1997). Activation mechanism of the MAP kinase ERK2 by dual phosphorylation. Cell 90, 859–869. 10.1016/s0092-8674(00)80351-7.

21. Lai, S., and Pelech, S. (2016). Regulatory roles of conserved phosphorylation sites in the activation T-loop of the MAP kinase ERK1. Mol Biol Cell 27, 1040–1050. 10.1091/mbc.E15-07-0527.

22. Nguyen, T.B., Manova, K., Capodieci, P., Lindon, C., Bottega, S., Wang, X.-Y., Refik-Rogers, J., Pines, J., Wolgemuth, D.J., and Koff, A. (2002). Characterization and expression of mammalian cyclin b3, a prepachytene meiotic cyclin. J Biol Chem 277, 41960–41969. 10.1074/jbc.M203951200.

23. Karasu, M.E., Bouftas, N., Keeney, S., and Wassmann, K. (2019). Cyclin B3 promotes anaphase I onset in oocyte meiosis. J Cell Biol 218, 1265–1281. 10.1083/jcb.201808091.

24. Bansal, S., and Tiwari, S. (2019). Mechanisms for the temporal regulation of substrate ubiquitination by the anaphase-promoting complex/cyclosome. Cell Div 14, 14. 10.1186/s13008-019-0057-5.

25. Chia, K.H., Takaki, H., Fujimitsu, K., Darling, S., Zou, J., Rappsilber, J., and Yamano, H. (2024). CDK1-PP2A-B55 interplay ensures cell cycle oscillation via Apc1-loop300. Cell Rep 43, 114155. 10.1016/j.celrep.2024.114155.

26. Fujimitsu, K., and Yamano, H. (2021). Dynamic regulation of mitotic ubiquitin ligase APC/C by coordinated Plx1 kinase and PP2A phosphatase action on a flexible Apc1 loop. The EMBO Journal 40, e107516. 10.15252/embj.2020107516.

27. Fujimitsu, K., Grimaldi, M., and Yamano, H. (2016). Cyclin-dependent kinase 1-dependent activation of APC/C ubiquitin ligase. Science 352, 1121–1124. 10.1126/science.aad3925.

28. Castro, A., Peter, M., Magnaghi-Jaulin, L., Vigneron, S., Galas, S., Lorca, T., and Labbé, J.C. (2001). Cyclin B/cdc2 induces c-Mos stability by direct phosphorylation in Xenopus oocytes. Mol Biol Cell 12, 2660–2671. 10.1091/mbc.12.9.2660.

29. Mendez, R., and Richter, J.D. (2001). Translational control by CPEB: a means to the end. Nat Rev Mol Cell Biol 2, 521–529. 10.1038/35080081.

30. Karasu, M.E., Bouftas, N., Keeney, S., and Wassmann, K. (2019). Cyclin B3 promotes anaphase I onset in oocyte meiosis. Journal of Cell Biology 218, 1265–1281. 10.1083/jcb.201808091.

31. Perez-Riverol, Y., Bai, J., Bandla, C., García-Seisdedos, D., Hewapathirana, S., Kamatchinathan, S., Kundu, D.J., Prakash, A., Frericks-Zipper, A., Eisenacher, M., et al. (2022). The PRIDE database resources in 2022: a hub for mass spectrometry-based proteomics evidences. Nucleic Acids Research 50, D543–D552. 10.1093/nar/gkab1038.

