## Supplementary Informations for "Greatwall depletion from Xenopus oocytes reveals a key role of the cyclin B/CDK1-PP2A-B55 balance in the coordination of meiotic events"

### Figure S1

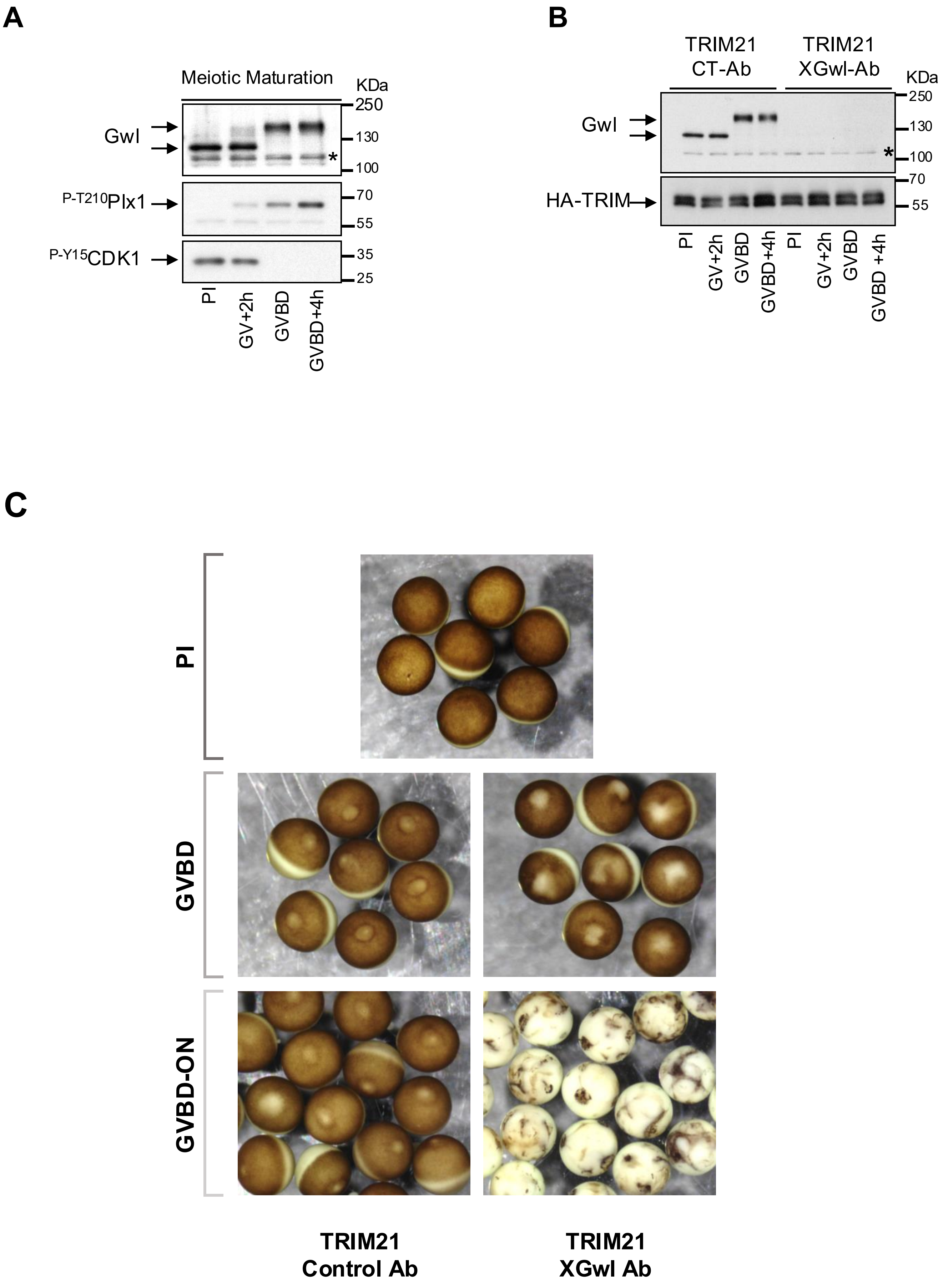
A
Meiotic Maturation
KDa
250
130
100
Gwl
*
P-T210Plx1
70
55
P-Y15CDK1
35
25
PI
GV+2h
GVBD
GVBD+4h
B
TRIM21
CT-Ab
TRIM21
XGwl-Ab
KDa
250
130
100
Gwl
*
70
HA-TRIM
55
PI
GV+2h
GVBD
GVBD+4h
PI
GV+2h
GVBD
GVBD +4h
C
PI
GVBD
GVBD-ON
TRIM21
Control Ab
TRIM21
XGwl Ab

**Figure S1. Effect of Gwl depletion in oocyte maturation. (A)** Prophase I arrested oocytes were treated with progesterone. At the indicated time-points oocytes were crushed by centrifugation and the cytoplasm recovered and supplemented with Laemmli solution. Immunoblot was performed at prophase I arrested oocytes (PI), two hours upon Pg addition (GV+2h), at Germinal Vesicle Breakdown (GVBD) and 4h upon GVBD, when oocytes were already arrested at Metaphase II (MII). Gwl, CDK1 phosphorylation on P-Y15 and Plx1 phosphorylation on P-T210 are shown **(B)** Prophase I arrested oocytes were injected with HA-TRIM21 mRNA and Control (anti-GST antibodies; CT-Ab) or Xenopus Gwl antibodies (Gwl-Ab). Sixteen hours later oocytes were treated with progesterone and used for immunoblotting at the specified time-points to examine Gwl and TRIM21 protein levels. **(C)** Prophase I arrested oocytes were injected with HA-TRIM21 mRNA and control or Xenopus Gwl antibodies. Sixteen hours later, oocytes were treated with progesterone and then monitored by video. Images were taken at prophase I, GVBD and 16h later.

Figure S2

A

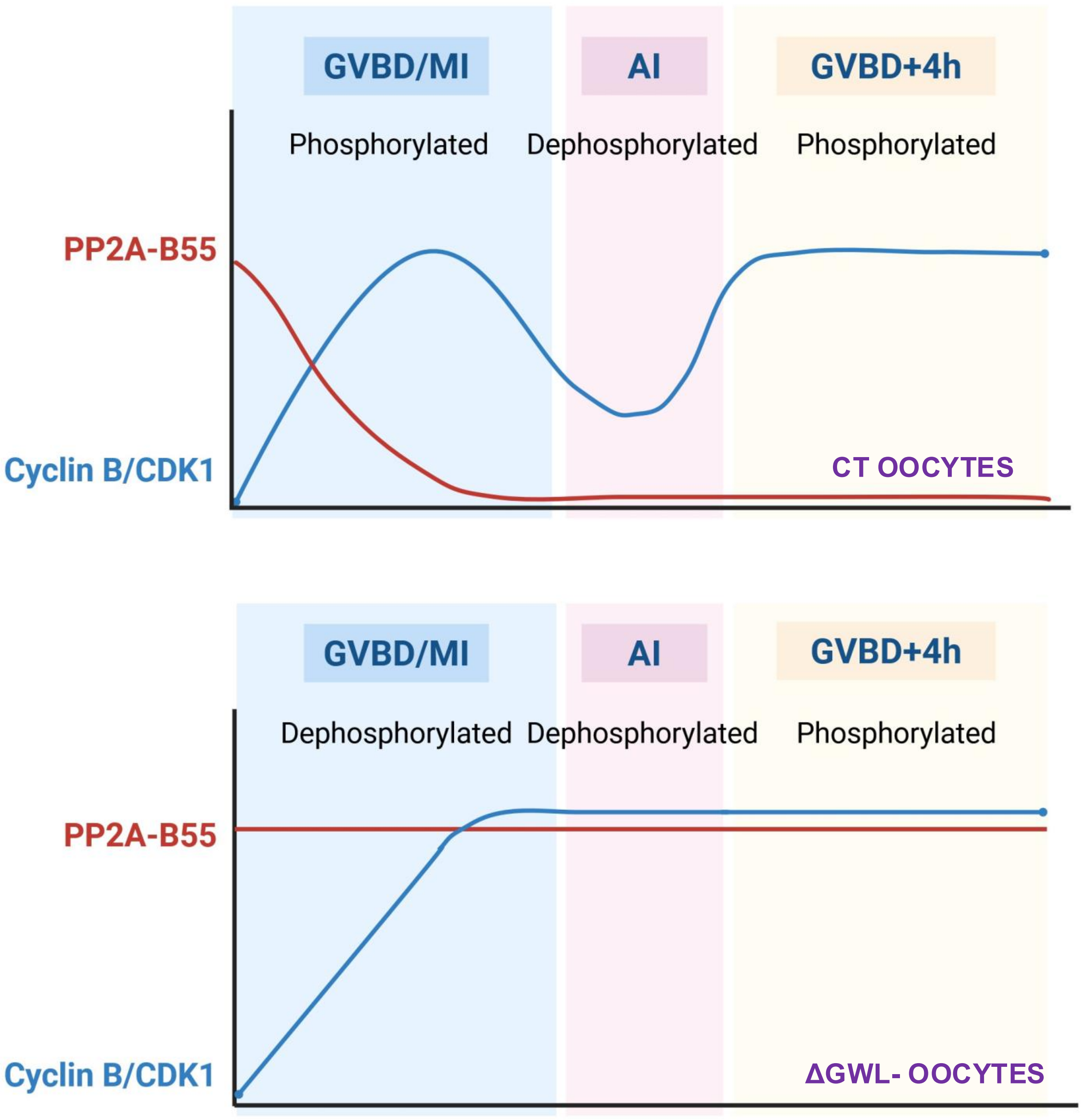
GVBD/MI
AI
GVBD+4h
Phosphorylated
Dephosphorylated
Phosphorylated
PP2A-B55
Cyclin B/CDK1
CT OOCYTES
GVBD/MI
AI
GVBD+4h
Dephosphorylated
Dephosphorylated
Phosphorylated
PP2A-B55
Cyclin B/CDK1
ΔGWL- OOCYTES

**Figure S2. Working model explaining the hypophosphorylation of meiotic substrates at GVBD and their hyperphosphorylation at GVBD+4H in Gwl depleted maturing oocytes.** Scheme representing the activities of PP2A-B55 and cyclin B/CDK1 in control and Greatwall depleted oocytes as well as the phosphorylation state of meiotic substrates at GVBD/ Metaphase I, Anaphase I and Metaphase II arrested oocytes (GVBD+4H). In Gwl depleted oocytes PP2A-B55 activity remains high throughout meiotic maturation due to the loss of Gwl and dephosphorylated Arpp19 and ENSA. Conversely, cyclin B/CDK1 activity abnormally increases due to the incapacity of the APC/C to degrade cyclin B. However, despite increased cyclin B/CDK1 activity, this kinase is unable to exceed PP2A-B55 activity at GVBD and substrates remain hypophosphorylated at this stage of meiosis in Gwl devoid oocytes. As oocytes progress in meiosis, cyclin B synthesis proceeds and cyclin B/CDK1 reach abnormal high activity that surpass the activity of the hyperactivated phosphatase promoting substrate hyperphosphorylation at GVBD+4h, a time point in which metaphase II is already established in control oocytes. Created with BioRender.com.
